# Targeting the mSWI/SNF Complex in POU2F-POU2AF Transcription Factor-Driven Malignancies

**DOI:** 10.1101/2024.01.22.576669

**Authors:** Tongchen He, Lanbo Xiao, Yuanyuan Qiao, Olaf Klingbeil, Eleanor Young, Xiaoli S. Wu, Rahul Mannan, Somnath Mahapatra, Sanjana Eyunni, Jean Ching-Yi Tien, Xiaoju Wang, Yang Zheng, NamHoon Kim, Heng Zheng, Siyu Hou, Fengyun Su, Stephanie J. Miner, Rohit Mehra, Xuhong Cao, Chandrasekhar Abbineni, Susanta Samajdar, Murali Ramachandra, Abhijit Parolia, Christopher R. Vakoc, Arul M. Chinnaiyan

**Affiliations:** Michigan Center for Translational Pathology, University of Michigan, Ann Arbor, MI, USA; Department of Pathology, University of Michigan, Ann Arbor, MI, USA; Department of Urology, Xiangya Hospital, Central South University, Changsha, Hunan, China; Rogel Cancer Center, University of Michigan, Ann Arbor, MI, USA; Cold Spring Harbor Laboratory, Cold Spring Harbor, NY, USA; Department of Biostatistics, School of Public Health, University of Michigan, Ann Arbor, MI, USA; Howard Hughes Medical Institute, University of Michigan, Ann Arbor, MI, USA; Aurigene Oncology Limited, Bangalore, India; Department of Urology, University of Michigan, Ann Arbor, MI, USA

**Keywords:** POU2F3, POU2AF1/2/3, mSWI/SNF complex, SMARCA2/4, proteolysis targeting chimera (PROTAC), small cell lung cancer (SCLC), multiple myeloma, IRF4

## Abstract

The POU2F3-POU2AF2/3 (OCA-T1/2) transcription factor complex is the master regulator of the tuft cell lineage and tuft cell-like small cell lung cancer (SCLC). Here, we found that the POU2F3 molecular subtype of SCLC (SCLC-P) exhibits an exquisite dependence on the activity of the mammalian switch/sucrose non-fermentable (mSWI/SNF) chromatin remodeling complex. SCLC-P cell lines were sensitive to nanomolar levels of a mSWI/SNF ATPase proteolysis targeting chimera (PROTAC) degrader when compared to other molecular subtypes of SCLC. POU2F3 and its cofactors were found to interact with components of the mSWI/SNF complex. The POU2F3 transcription factor complex was evicted from chromatin upon mSWI/SNF ATPase degradation, leading to attenuation of downstream oncogenic signaling in SCLC-P cells. A novel, orally bioavailable mSWI/SNF ATPase PROTAC degrader, AU-24118, demonstrated preferential efficacy in the SCLC-P relative to the SCLC-A subtype and significantly decreased tumor growth in preclinical models. AU-24118 did not alter normal tuft cell numbers in lung or colon, nor did it exhibit toxicity in mice. B cell malignancies which displayed a dependency on the POU2F1/2 cofactor, POU2AF1 (OCA-B), were also remarkably sensitive to mSWI/SNF ATPase degradation. Mechanistically, mSWI/SNF ATPase degrader treatment in multiple myeloma cells compacted chromatin, dislodged POU2AF1 and IRF4, and decreased IRF4 signaling. In a POU2AF1-dependent, disseminated murine model of multiple myeloma, AU-24118 enhanced survival compared to pomalidomide, an approved treatment for multiple myeloma. Taken together, our studies suggest that POU2F-POU2AF-driven malignancies have an intrinsic dependence on the mSWI/SNF complex, representing a therapeutic vulnerability.

## Introduction

Small cell lung cancer (SCLC) is an aggressive, fast-evolving subtype of lung cancer with a high growth rate and early metastasis propensity, often resulting in a more advanced disease stage at diagnosis^1,2^. Consequently, the overall prognosis for SCLC is generally poorer compared to non-small cell lung cancer (NSCLC)^3^. Unlike NSCLC, where substantial progress has been achieved with immune checkpoint blockade therapies, effective targeted therapies for SCLC remain elusive^4^. Comprehensive genome sequencing of SCLC tumors has revealed a high mutational load in this disease, with most tumors possessing inactivating mutations or deletions of *RB1* and *TP53*, but few actionable targets have been identified^5^. Thus, there is an urgent need for innovative therapeutic strategies that address the distinct biology of SCLC and enhance patient outcomes.

Prior analysis of human SCLC tumors revealed that SCLC could be characterized by the expression pattern of certain transcription factors (TFs) or transcriptional regulators, including ASCL1 (achaete-scute family bHLH transcription factor 1), NeuroD1 (neurogenic differentiation factor 1), POU2F3 (POU domain class 2 transcription factor 3; also known as OCT-11), and YAP1 (yes-associated protein 1), exemplifying SCLC as a TF-driven malignancy^6–9^. ASCL1-driven SCLC (SCLC-A) and NeuroD1-driven SCLC (SCLC-N) manifest a neuroendocrine phenotype, while POU2F3-driven SCLC (SCLC-P) is characterized as a tuft cell-like variant^9^. Prior studies revealed that POU domain class 2 TFs uniquely rely on coactivators to achieve their lineage-defining functions in B cells^10–13^. More recently, in tuft cell-like SCLC cells, the coactivators of POU2F3 (POU2AF2 and POU2AF3) were found to endow POU2F3 with a critical transactivation domain by forming a master regulator complex, which supports enhancer-mediated cancer-promoting gene activation in SCLC-P cells^14–16^. This indicates a potential therapeutic vulnerability in patients with tuft cell-like SCLC whereby strategies aimed at blocking POU2F3-POU2AF2/3 function may lead to clinical benefit.

The mammalian switch/sucrose non-fermentable (mSWI/SNF) chromatin remodeling complex acts as a pivotal regulator of gene expression and chromatin architecture, thereby orchestrating fundamental cellular processes crucial for homeostasis and development^17^. The ATPase subunit of this complex harnesses energy from ATP hydrolysis to reposition or eject nucleosomes at non-coding regulatory elements, facilitating unobstructed DNA access for the transcriptional machinery^18–20^. Recent investigations have elucidated alterations in the genes encoding constituent subunits of the mSWI/SNF complex in over 25% of human malignancies^21,22^. Our group recently discovered that androgen receptor (AR)-driven prostate cancer cells are preferentially dependent on the chromatin remodeling function of the mSWI/SNF complex^23^. We identified a novel mSWI/SNF ATPase proteolysis targeting chimera (PROTAC) degrader that dislodges AR and its cofactors from chromatin, disabling their core enhancer circuitry and attenuating downstream oncogenic gene programs^23^. Similar observations have been reported in other TF-driven malignancies like acute myeloid leukemia^24,25^, highlighting the broad applicability of targeting the mSWI/SNF complex in a variety of malignancies.

In this study, we identified an enhanced dependency on the mSWI/SNF complex in POU2F3-driven SCLC cells through CRISPR screening and pharmacological validation. Epigenomics analyses revealed that inactivation of the mSWI/SNF complex preferentially obstructed chromatin accessibility of POU2F3 complexes, leading to a dramatic downregulation of POU2F3 signaling. Critically, treatment with an orally bioavailable mSWI/SNF ATPase PROTAC degrader resulted in significant tumor growth inhibition in preclinical models of POU2F3-driven SCLC without significant effects in other subtypes of SCLC xenografts. Furthermore, our investigations extended to other POU2AF1 complex-dependent B cell malignancies, including multiple myeloma, wherein sensitivity to the mSWI/SNF ATPase PROTAC degrader was observed *in vitro* and *in vivo*. These findings collectively show the potential of targeting the mSWI/SNF complex in POU2F-POU2AF-driven malignancies and suggest that development of mSWI/SNF degraders should be pursued as targeted therapies for patients with these types of cancers.

## Results

### Dependence of SCLC-P Cells on the mSWI/SNF Complex

SCLCs are genetically driven by loss of function (LOF) alterations in tumor suppressor genes *RB1* and *TP53*^5^, with distinct expression patterns of certain TFs or transcriptional regulators leading to four molecular subtypes (SCLC-A, SCLC-N, SCLC-P, and SCLC-Y (YAP1))^6^. Functional genomics analyses have underscored the critical roles of these TFs or coactivators in each SCLC molecular subtype. However, unlike kinases, many TFs have been perceived as undruggable targets due to their enrichment of intrinsically disordered regions within their structures, indicating potential challenges in devising ASCL1 or POU2F3-direct targeting strategies. Considering this, we hypothesized that druggable targets selective to SCLC subtypes could be identified via a loss-of-function CRISPR/Cas9 screen. Accordingly, we conducted a functional domain-targeted CRISPR/Cas9 screen co-targeting paralog pairs of kinases, phosphatases, epigenetic regulators, and DNA binding proteins in three SCLC-A and three SCLC-P cell lines (**Fig. 1A**). Dependency scores (beta scores) for 4,341 single-gene and 4,387 double-gene knockouts were calculated using MAGeCK^26^. Comparing beta scores between SCLC-A and SCLC-P cell lines, we observed dramatic dependency differences for lineage TFs ASCL-1 and POU2F3. Surprisingly, we also identified a strong dependency bias of multiple components of the mSWI/SNF complex in SCLC-P cells (**Fig. 1B-D, Fig. S1A-B, Supplementary Table 1**).

**Figure 1.**
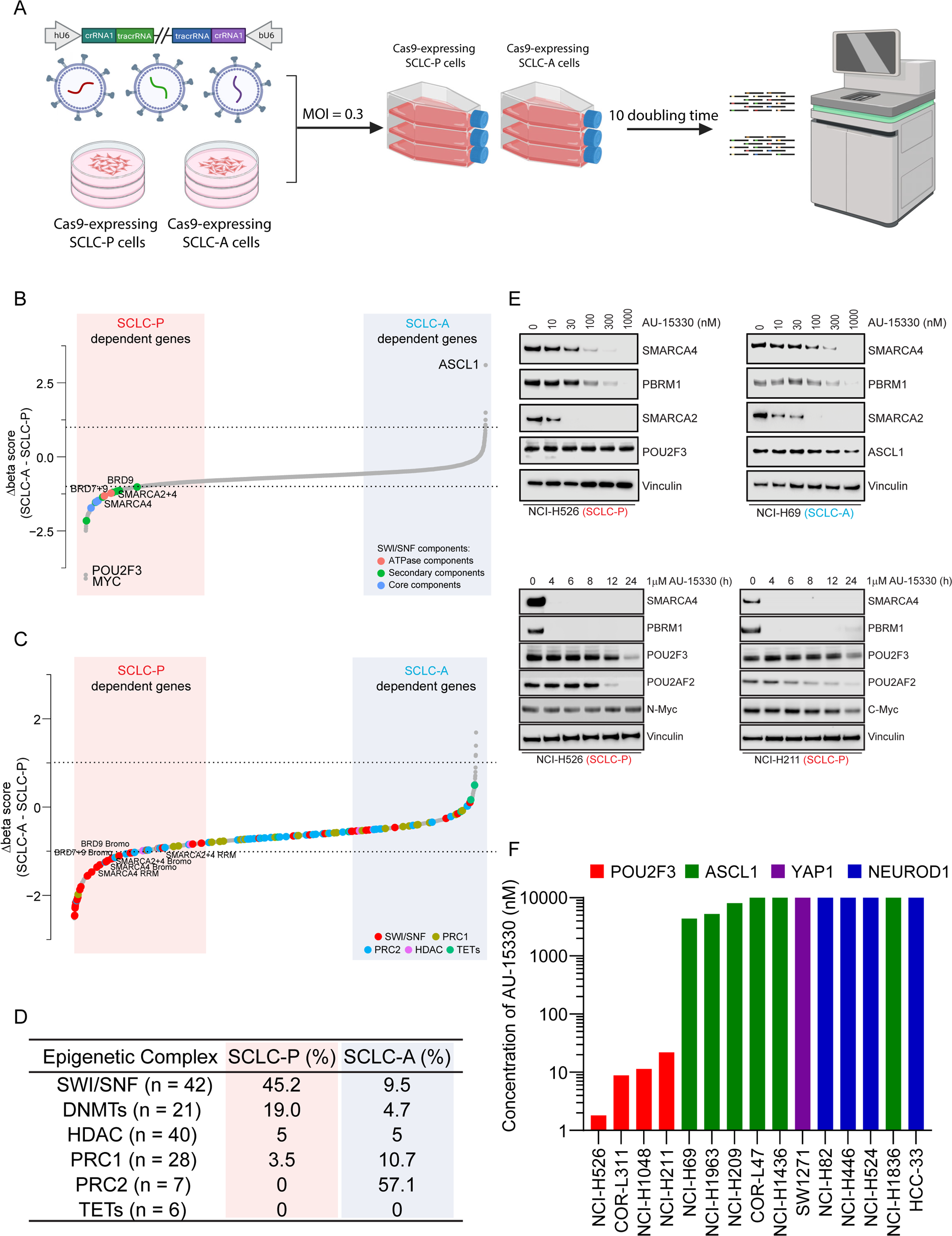
Dependence of SCLC-P cells on the mSWI/SNF complex. (A) A schematic representation of the dual-sgRNA, domain-focused CRISPR screening designed to identify druggable epigenetic targets selective for SCLC subtypes. (B) Beta scores pertaining to all CRISPR screen targeted genes across both SCLC-P and SCLC-A cell lines (n = 5308). (C) Beta scores highlighting epigenetic regulators in SCLC-P and SCLC-A cell lines (n = 3292). (D) Percentage of different epigenetic complexes in SCLC-P and SCLC-A cell lines (top 10% for each). PRC1, polycomb repressive complex 1; PRC2, polycomb repressive complex 2; HDAC, histone deacetylase; TET, ten-eleven translocation family proteins. (E) Immunoblot analysis of indicated proteins in SCLC-P and SCLC-A cells post-treatment with varying time points or concentrations of AU-15330. Vinculin serves as the control for protein loading in all immunoblots. (F) Compilation of the IC_50_ values for AU-15330 in SCLC cell lines representing four molecular subtypes.

We hypothesized that this selective dependency might originate from a POU2F3-imposed requirement on the mSWI/SNF complex. Among the mSWI/SNF complex components, only ATPases and bromodomain containing 9 (BRD9) were found to be directly targetable by recently developed PROTAC degraders, which have been engineered to induce target protein degradation through the ubiquitin-proteasome system (**Fig. S1B, Supplementary Table 1**)^27,28^. Our team recently showcased the promising anti-tumor efficacy of the PROTAC degrader targeting the mSWI/SNF ATPase subunit in preclinical models of AR-driven prostate cancer^23^. Here, we evaluated the efficacy of this mSWI/SNF ATPase PROTAC degrader, AU-15330, across a spectrum of SCLC cell lines. AU-15330 treatment resulted in time and dose-dependent degradation of mSWI/SNF ATPases (SMARCA2 and SMARCA4) and PBRM1 in cell lines encompassing all four molecular subtypes of SCLC (**Fig. 1E, Fig. S1C**). Protein levels of POU2F3 and its coactivator POU2AF2 were also decreased in SCLC-P cells treated with AU-15330 at extended time points (12 and 24 hours, **Fig. 1E, Fig. S1C**). Despite degradation of target mSWI/SNF ATPase proteins across subtypes, AU-15330 exhibited a preferential growth inhibitory effect in SCLC-P cells compared to all non-POU2F3 SCLC cell line models (**Fig. 1F, Fig. S1D-E**). Taken together, our functional CRISPR/Cas9 screen, complemented by secondary pharmacological validation, pinpointed the mSWI/SNF complex and its catalytic ATPase subunit as novel epigenetic dependencies in SCLC-P cells.

### Mechanism of Action of mSWI/SNF Complex Inactivation in SCLC-P Cells

Experiments were next performed to elucidate the mechanism of action underlying the selective growth inhibitory effects of the mSWI/SNF ATPase PROTAC degrader in SCLC-P cells. Given the primary role of the mSWI/SNF complex in modulating chromatin accessibility by altering nucleosome positioning along DNA, we employed Assay for Transposase-Accessible Chromatin using sequencing (ATAC-seq) in SCLC-P and SCLC-A cells post AU-15330 treatment. As depicted in **Fig. 2A** and **Fig. S2A**, four-hour treatment with AU-15330 triggered rapid and genome-wide chromatin compaction at regulatory regions in both SCLC-P and SCLC-A cells. *De novo* motif analysis of the AU-15330-compacted sites revealed that POU motif-containing sites were predominantly affected across the genome in SCLC-P cells (**Fig. 2B, Fig. S2B-F**). Conversely, the ASCL1 motif was only mildly impacted upon AU-15330 treatment in ASCL1-expressing NCI-H69 cells (**Fig. 2B, Fig. S2G-H**), suggesting that chromatin accessibility of ASCL1-targeting regions is largely independent of the mSWI/SNF complex. Concurrent with the loss of chromatin accessibility, chromatin immunoprecipitation followed by sequencing (ChIP-seq) showed diminished chromatin binding of POU2F3 and its coactivators (POU2AF2, POU2AF3) at the AU-15330-mediated compacted sites, as examined by tagging endogenous or exogenous POU2F3 and its coactivators in SCLC-P cell lines (**Fig. 2C, Fig. S3A-F**).

**Figure 2.**
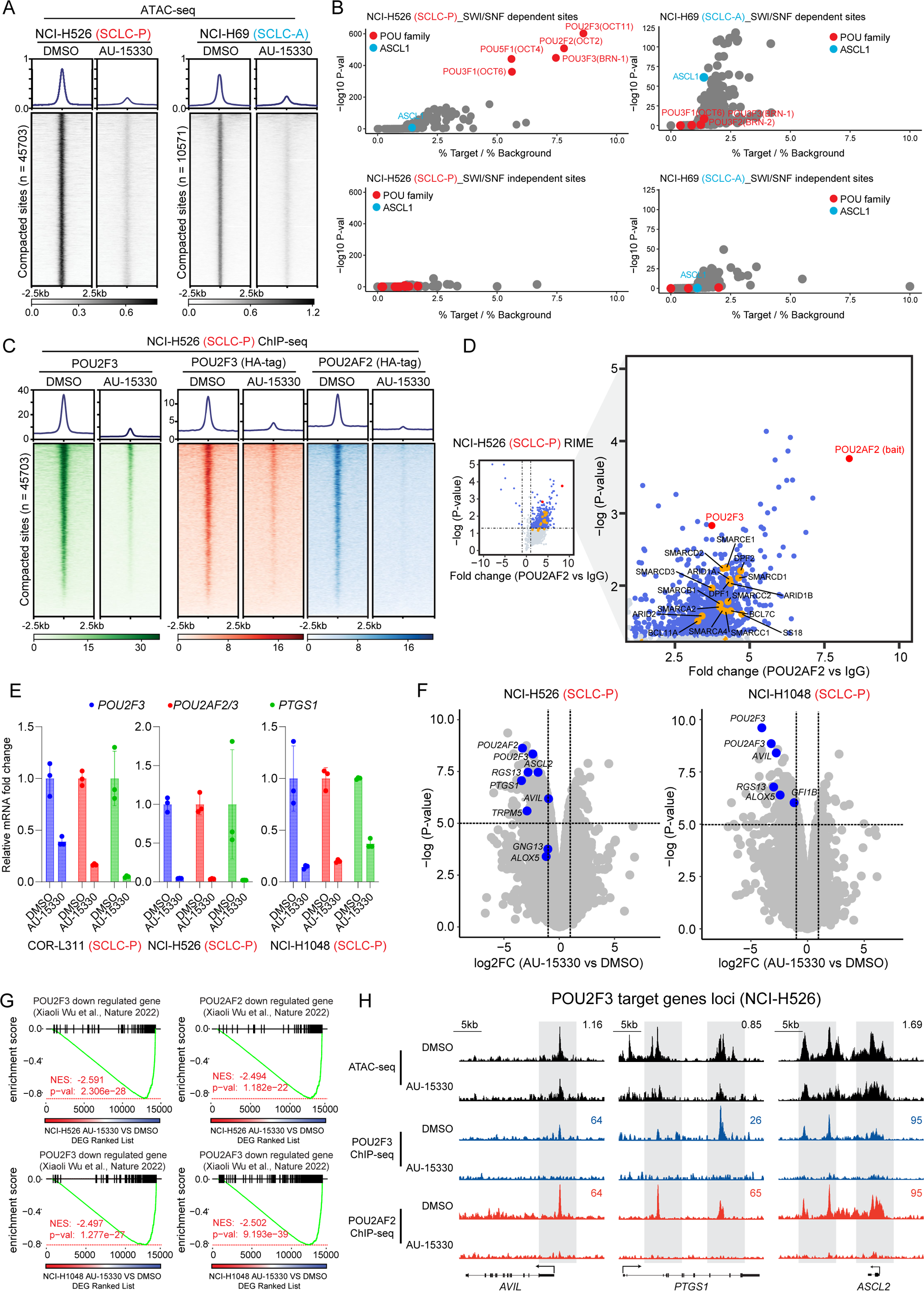
The POU2F3 transcription factor complex is evicted from chromatin in SCLC-P cells upon mSWI/SNF ATPase degradation. (A) Chromatin compaction induced by mSWI/SNF ATPase degradation. Visualization of ATAC-seq read-density in NCI-H526 (SCLC-P) and NCI-H69 (SCLC-A) cells post-treatment for 4 hrs with either vehicle or 1 mM AU-15330. (B) Analysis of fold change and significance level for HOMER motifs that are enriched within sites dependent and independent of the mSWI/SNF complex in NCI-H526 and NCI-H69 cells. (C) ChIP–seq read-density heatmaps representing POU2F3 (green), HA-POU2F3 (red), and HA-POU2AF2 (blue) at AU-15330-compacted genomic sites in NCI-H526 cells following treatment with DMSO or AU-15330. (D) Volcano plot detailing proteins that interact with POU2AF2, as identified by POU2AF2 RIME analysis in NCI-H526 cells. mSWI/SNF components highlighted in orange. (E) Expression levels of *POU2F3*, *POU2AF2/3*, and *PTGS1* as assessed by QPCR in the indicated cell lines after being treated with vehicle or AU-15330. (F) Volcano plot visualizing the overall transcriptomic alterations as assessed by RNA-seq in NCI-H526 and NCI-H1048 cells post-treatment with vehicle or AU-15330. Canonical POU2F3 target genes are highlighted in blue. (G) GSEA plots illustrating genes regulated by POU2F3 and its coactivators POU2AF2 and POU2AF3. The plots employ a gene signature ranked by fold change in AU-15330-treated NCI-H526 and NCI-1048 cells. DEG, differentially expressed gene. (H) Combined ATAC-seq and ChIP-seq tracks for *AVIL*, *PTGS1*, and *ASCL2* in NCI-H526 with and without AU-15330 treatment.

Given the pronounced impact on POU motif-containing sites upon mSWI/SNF complex inactivation, we hypothesized an association between the mSWI/SNF complex and the POU2F3 complex in SCLC-P cells. To explore this, we conducted Fast Protein Liquid Chromatography (FPLC) experiments to size fractionate the nuclear lysate from two SCLC-P cell lines. We observed several mSWI/SNF complex components (SMARCD1, ARID1A, and SS18), POU2F3, and POU2AF2 co-expressed in the large nuclear fractions (**Fig. S4A**), suggesting a potential coexistence of the POU2F3 complex and the mSWI/SNF complex within a large nuclear protein complex. Further, Rapid Immunoprecipitation Mass Spectrometry of Endogenous Proteins (RIME) analysis of POU2F3 and its coactivators’ interactome revealed multiple key mSWI/SNF components coimmunoprecipitated with POU2F3 and its coactivators (**Fig. 2D, Fig. S4B-E, Supplementary Table 1**), affirming the physical interaction between the POU2F3 complex and the mSWI/SNF complex in SCLC-P cells. Real-time quantitative reverse transcription PCR (qRT-PCR) and global transcriptomic profiling via RNA sequencing (RNA-seq) showcased significant downregulation of *POU2F3*, *POU2AF2/3*, and their downstream targets (e.g., *PTGS1*) in multiple SCLC-P cell lines (**Fig. 2E-F**). The Gene Set Enrichment Analysis (GSEA) of global AU-15330-mediated transcriptomic alterations reflected a high concordance between mSWI/SNF inactivating gene signatures and transcriptional signatures associated with genetic knockout of *POU2F3* and its coactivators (**Fig. 2G-H, Fig. S4F**)^15^. Collectively, our multi-omics analysis suggests that the POU2F3 complex necessitates the mSWI/SNF complex to modulate chromatin accessibility at its DNA binding regions, thereby transactivating the POU2F3 downstream signaling pathway in SCLC-P cells.

### Selective Inhibition of SCLC-P Xenograft Tumor Growth by AU-24118

To enhance the translational potential of our findings, we developed a first-in-class, orally bioavailable SMARCA2/4 PROTAC degrader, named AU-24118, as AU-15330 does not possess optimal oral bioavailability. AU-24118 inhibited growth of SCLC-P cell lines at nanomolar levels compared to SCLC-A, SCLC-N, and SCLC-Y cell lines (**Fig. S5A**), corroborating results observed with AU-15330 that the SCLC-P molecular subtype is preferentially sensitive to mSWI/SNF inactivation (**Fig. S1D**).

AU-24118 was administered at 15 mg/kg by oral gavage (o.g.), three times weekly, to immunodeficient mice bearing subcutaneous SCLC tumors representing the SCLC-P (NCI-H526, NCI-H1048) and SCLC-A (NCI-H69) molecular subtypes (**Fig. 3A**). Notably, significant reductions in SCLC-P tumor volumes (**Fig. 3B**) and tumor weights (**Fig. S5B**) were observed post-oral administration of AU-24118, with no discernible changes in body weights (**Fig. S6A**). Conversely, AU-24118 treatment did not significantly alter tumor growth of NCI-H69 SCLC-A xenografts (**Fig. 3B, Fig. S5B, Fig. S6A**), thereby confirming the selective anti-tumor efficacy of mSWI/SNF ATPase degraders in SCLC-P preclinical models. Aligning with our *in vitro* observations, SCLC-P tumors treated with AU-24118 exhibited significant degradation of its direct targets (SMARCA2/4 and PBRM1), which ensued in downregulation of POU2F3, POU2F3 coactivators, and downstream target expression (GFI1B) (**Fig. 3C**). Levels of cleaved PARP were also increased in SCLC-P tumors treated with AU-24118, while N-MYC levels decreased with treatment (**Fig. 3C**). Further, histopathological assessments performed on AU-24118-treated SCLC-P tumors showed increased apoptotic bodies and intra-tumoral nuclear and necrotic debris in contrast to highly cellular and monotonous appearing, high-grade vehicle-treated tumor samples (**Fig. 3D, Fig. S5C**). This was corroborated by fluorometric terminal deoxynucleotidyl transferase (TUNEL) assay analysis showing a significant increase in TUNEL-positive cells in SCLC-P but not SCLC-A tumors (**Fig. 3E**). Additionally, immunohistochemistry (IHC) confirmed a dramatic loss of SMARCA4 and POU2F3 protein expression in the AU-24118-treated SCLC-P samples, as well as decreased DCLK1 expression - a tuft cell marker (**Fig. 3D, Fig. S5C**). Despite no changes in tumor growth in the SCLC-A xenografts, western blot and IHC analysis of tumors confirmed on-target drug activity of AU-24118 as indicated by efficient loss of SMARCA4, SMARCA2, and PBRM1 (**Fig. 3C, Fig. S5D**).

**Figure 3.**
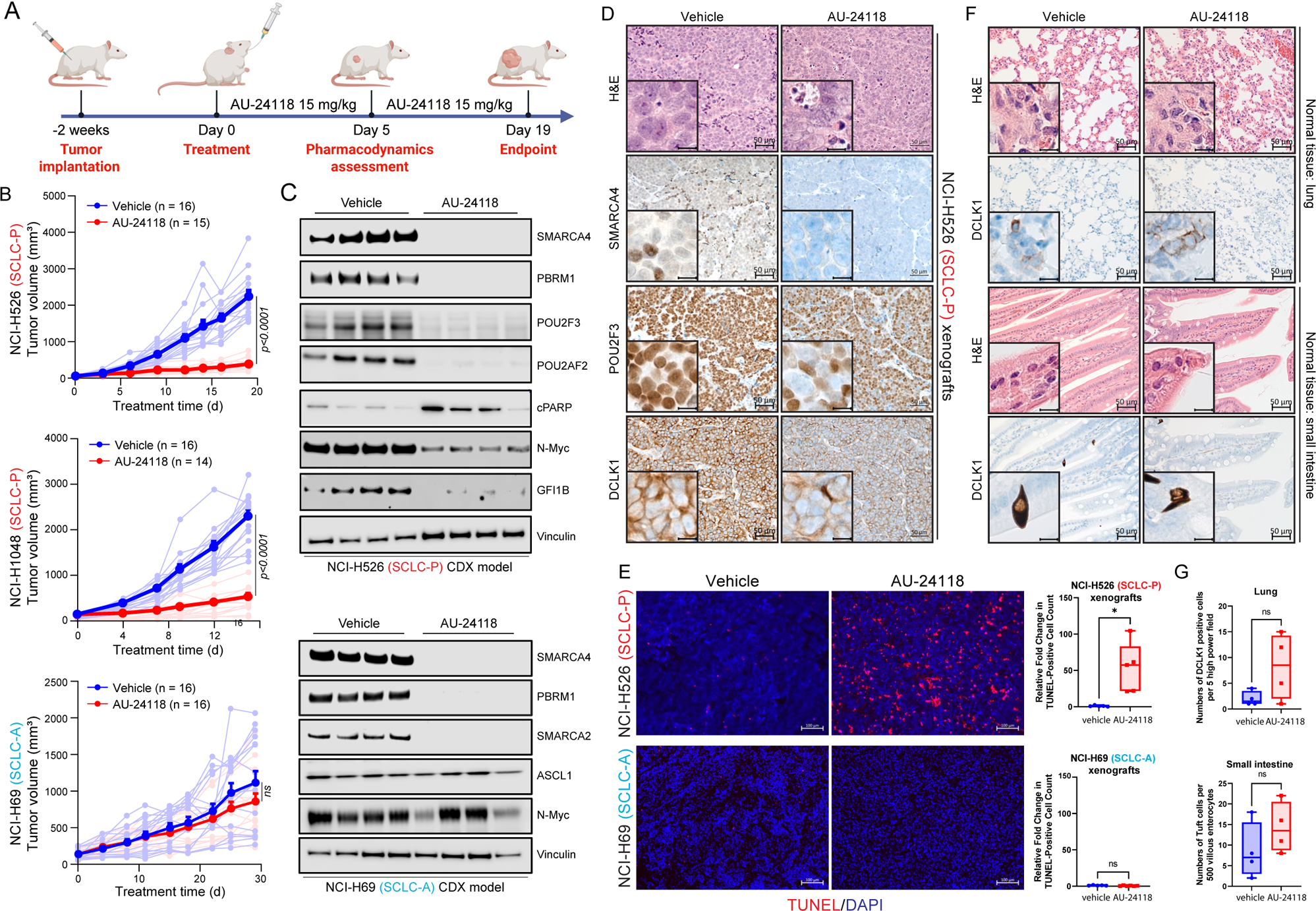
Selective inhibition of SCLC-P xenograft tumor models employing an orally bioavailable mSWI/SNF ATPase degrader. (A) Overview of the AU-24118 efficacy study conducted using SCLC xenograft models. (B) Analysis of tumor volume in indicated SCLC xenograft models upon treatment with AU-24118, measured bi-weekly using calipers (analyzed with a two-way ANOVA). (C) Immunoblot illustrating levels of the indicated proteins in SCLC-P and SCLC-A xenografts after 5 days of AU-24118 administration. Vinculin is utilized as the loading control across immunoblots. CDX, cell line-derived xenograft. (D) Representative H&E staining with corresponding IHC analyses for SMARCA4, POU2F3, and DCLK1 after 5 days of treatment with AU-24118 in NCI-H526 xenografts (scale=50mm). The inset scale=20mm. (E) (Left) Representative DAPI and TUNEL staining from xenografts from indicated cell lines after 5 days of AU-24118 treatment (scale=100mm). (Right) Quantitative evaluation of TUNEL staining of respective SCLC xenografts for 5 days. T-tests were used to calculate the significance. P value < 0.05 in the top panel. (F) Representative H&E staining of murine lung and small intestine with corresponding tuft cell marker DCLK1 IHC after *in vivo* administration of AU-24118 at study endpoint (scale=50mm). Magnified views of intestinal enterocytes and lung alveolar epithelium in H&E and corresponding DCLK1 IHC shown in insets (scale=20mm). (G) DCLK1 cell positivity in lung and small intestine for endpoint evaluation. AU-24118 (15mg/kg) dosed. Ns, not significant (t-tests).

A comprehensive and detailed histopathological assessment was performed to identify any effects of AU-24118 on the lung, liver, spleen, kidney, small intestine, and colon of treated mice. With brightfield microscopy, no remarkable changes or toxic effects were noted with AU-24118 as compared to vehicle-treated animals (**Fig. 3F, Fig. S6B**). As validation, we performed IHC for a canonical chemosensory marker, DCLK1, in small intestine and lung tissues of the AU-24118 and vehicle-treated mice to rule out toxicity at the cyto-molecular level. Microscopic evaluation followed by quantification did not show any statistically significant increase in levels of DCLK1 between vehicle and AU-24118-treated samples (**Fig. 3F-G**). Collectively, these results position AU-24118 as the first orally bioavailable mSWI/SNF ATPase degrader with potent anti-tumor efficacy and no signs of toxicity in preclinical models of the SCLC-P molecular subtype.

### POU2AF1-Dependent B Cell Malignancies Exhibit Sensitivity to SMARCA2/4 PROTAC Degraders

POU2AF1 (OCA-B) is a coactivator that interfaces with TFs POU2F1 (OCT1) and POU2F2 (OCT2) at octamer motifs, orchestrating B cell development, maturation, and germinal center formation^13,29–31^. In diffuse large B cell lymphoma (DLBCL) cells, the *POU2AF1* locus is the most BRD4-overloaded super-enhancer, with subsequent analyses underscoring the importance of POU2AF1 to DLBCL growth and other B cell malignancies^32^. Prior investigations also highlighted homology between POU2AF1 and POU2F3 coactivators (POU2AF2, POU2AF3), with all three genes residing within the same genomic loci^15^, suggesting potential overlapping coactivator functions. Using data from the DepMap Project^33,34^, we discovered that POU2AF1 is indispensable for the growth of DLBCL and multiple myeloma (MM) cells but is non-essential for other cancer types (**Fig. 4A, Fig. S7A**).

**Figure 4.**
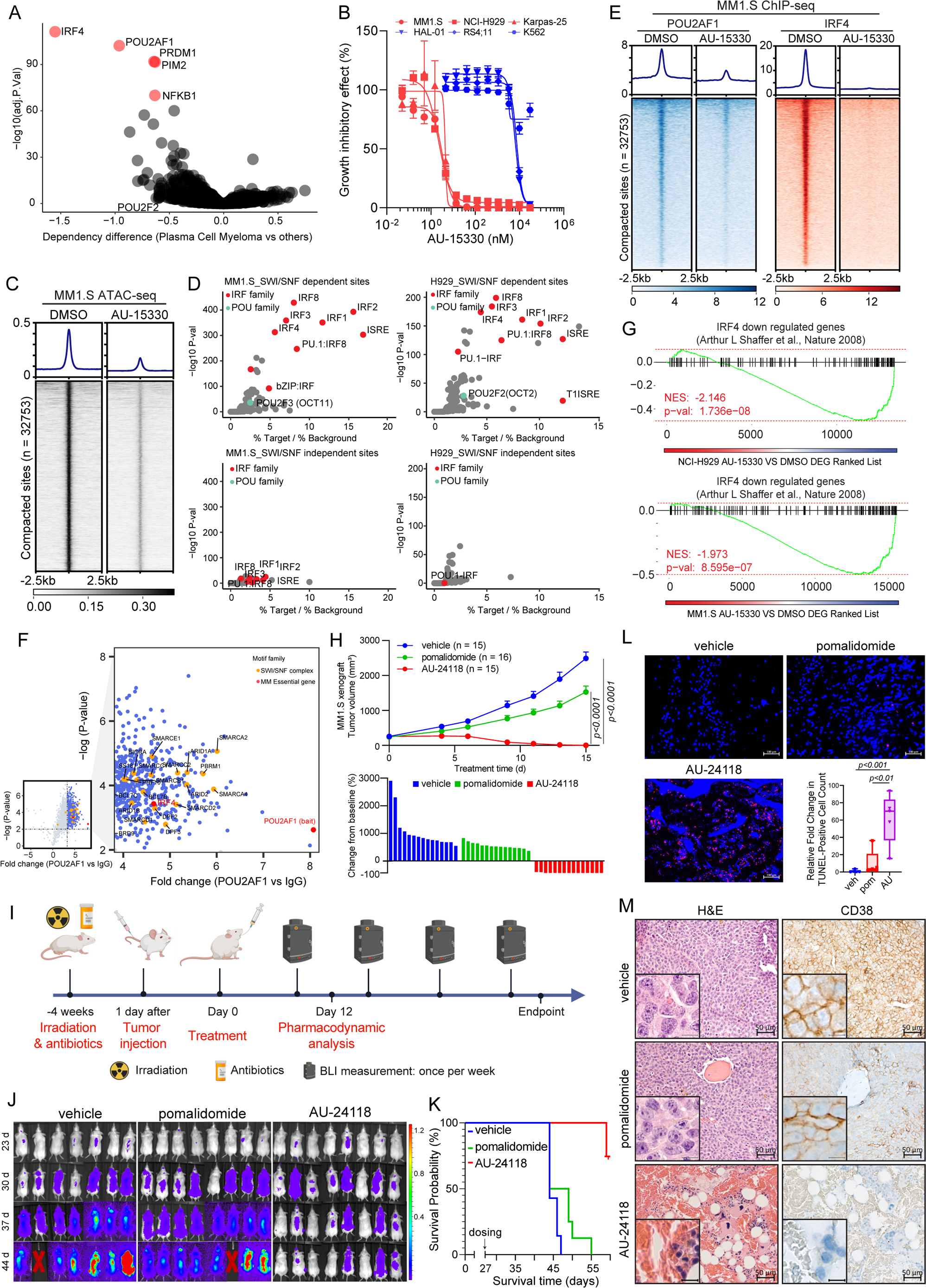
POU2AF1-driven B cell malignancies are dependent on the mSWI/SNF complex. (A) Scatter plot depicting gene dependency difference of all plasma cell myeloma versus other cancer types based on DepMap. The red circles indicate the top 5 essential genes among others. (B) Representative hematological cancer cell lines showing dose-response curves of AU-15330 at varying concentrations for five days. Sensitive cell lines are in red while relatively resistant cell lines in blue. (C) ATAC-seq read-density heatmaps from MM1.S cells treated with DMSO or 1 mM AU-15330 for 4 hours (n = 2 biological replicates). (D) Analysis of fold change and significance level for HOMER motifs that are enriched within sites dependent and independent of the mSWI/SNF complex after 4 hrs AU-15330 treatment in MM1.S cells (left panels) and NCI-H929 cells (right panels). (E) ChIP-seq read-density heat maps for POU2AF1 and IRF4 at the AU-15330-compacted genomic sites in MM1.S cells after treatment with DMSO or AU-15330 (1μM) for 6 hrs. (F) Volcano plot detailing proteins that interact with POU2AF1, as identified by POU2AF1 RIME analysis in MM1.S cells. mSWI/SNF components highlighted in orange. (G) GSEA plots illustrating genes regulated by IRF4. The plots use a gene signature ranked by fold change from AU-15330 treated NCI-H929 (top) and MM1.S (bottom) cells. (H) (Top) Analysis of tumor volumes in the MM1.S xenograft model upon treatment with AU-24118 and pomalidomide, measured bi-weekly using calipers (analyzed with a two-way ANOVA). (Bottom) Waterfall plot of tumor volumes at endpoint. (I) Overview of the MM1.S multiple myeloma disseminated xenograft model efficacy study. (J) Bioluminescence of images of MM1.S disseminated xenograft model after different treatments. The mice were monitored once per week. The signal intensity of bioluminescence represented the tumor burden (x10^8^ photons/sec/cm^2^/steradian). Pomalidomide (10mg/kg) and AU-24118 (15mg/kg) dosed. (K) Kaplan-Meier survival curve of MM1.S disseminated xenograft model after pomalidomide (10mg/kg) and AU-24118 (15mg/kg) treatment. (L) Representative DAPI and TUNEL staining from the MM1.S disseminated xenograft model and quantitative evaluation from TUNEL staining for pomalidomide (10mg/kg) and AU-24118 (15mg/kg) treatment for 12 days. (M) Representative H&E and CD38 IHC staining of spinal vertebral marrow after *in vivo* administration of pomalidomide (10mg/kg) and AU-24118 (15mg/kg) for 12 days.

Given the selective dependency of SCLC-P cells on the mSWI/SNF complex, we assessed whether POU2AF1-dependent B cell malignancies also exhibited sensitivity to SMARCA2/4 degraders. Three initial MM cell lines tested showed enhanced sensitivity to growth inhibition by AU-15330 compared to three cell lines from other hematological malignancies (**Fig. 4B**). The MM cell lines showed rapid loss of target proteins (SMARCA4, PBRM1) as well as POU2AF1 and c-MYC at extended time points (**Fig. S7B**). Determination of cell viability across an expanded panel of MM and DLBCL cell lines found that seven out of ten MM cell lines and 7 out of 17 DLBCL cell lines exhibited IC_50_ values below 200 nM for AU-15330 (**Fig. S7C-D**). These data indicated that a subset of POU2AF1-dependent MM and DLBCL cells exhibit an enhanced dependency on the mSWI/SNF complex.

Experiments were undertaken to define the mechanism of action of mSWI/SNF ATPase degraders in sensitive MM cell lines. Chromatin accessibility changes in two AU-15330 sensitive MM cell lines (MM.1S and NCI-H929) were assessed through ATAC-seq. As observed in SCLC-P cells, AU-15330 induced genome-wide chromatin compaction in both MM cell lines tested (**Fig. 4C, Fig. S8A**). Interestingly, *de novo* motif analysis of AU-15330-compacted sites revealed that, unlike SCLC-P cells, interferon regulatory factor (IRF) motif-containing sites, rather than POU motif-containing sites, were most enriched in MM cells (**Fig. 4D, Fig. S8B-E**). Given IRF4’s central role in MM tumorigenesis^35^ and the absence of POU motifs in the MM ATAC-seq data, we postulated that POU2AF1 might act as a novel transcriptional coactivator for IRF4 in MM cells. Analysis of the DepMap data revealed a significant positive correlation between the essentiality scores of IRF4 and POU2AF1 in MM cells, whereas sole knockout of POU2F1 and POU2F2 were less essential (**Fig. S8F**). Subsequent ChIP-seq analysis revealed a notable loss of POU2AF1 and IRF4 binding within AU-15330-compacted sites. Strikingly, *de novo* motif analysis revealed significant enrichment of IRF motifs within POU2AF1 binding sites, suggesting potential formation of a complex containing these regulators at certain genomic loci, such as enhancers near the *MYC* gene (**Fig. 4E, Fig. S8G-I**). Moreover, a RIME experiment conducted to delineate the interactome of POU2AF1 in MM.1S cells revealed numerous mSWI/SNF complex components co-immunoprecipitated with POU2AF1 (**Fig. 4F**). A robust association between IRF4 and POU2AF1 was also discovered in MM.1S cells (**Fig. 4F**). Global transcriptomic profiling via RNA-seq showcased significant downregulation of IRF4 downstream targets^35^ in two MM cell lines treated with AU-15330 (**Fig. 4G**).

To evaluate the therapeutic potential of targeting the mSWI/SNF ATPases in MM, we investigated the anti-tumor efficacy of the oral degrader, AU-24118, across diverse MM preclinical models. Initially, immunodeficient mice bearing MM subcutaneous tumors (MM.1S, NCI-H929, and Karpas-25) were treated with either vehicle, pomalidomide (10 mg/kg, o.g., five times weekly), carfilzomib (5 mg/kg, i.v., bi-weekly), or AU-24118 (15 mg/kg, o.g., three times weekly) (**Fig. S9A**). In all three models, AU-24118 significantly decreased tumor volumes and weights compared to pomalidomide or carfilzomib, without notable alterations in body weights (**Fig. 4H**, **Fig. S9B-E**). Notably, in the AU-24118 treatment arm, tumor regression was observed in all animals with no palpable tumors at endpoint (bottom panel, **Fig. 4H**). Western blot analysis of tumors from the MM1.S model showed marked loss of target proteins (SMARCA2, SMARCA4, PBRM1) as well as c-MYC and POU2AF1 (**Fig. S9F**). A disseminated orthotopic xenograft model of MM was next used to more physiologically recapitulate the disease state in patients. Luciferase and green fluorescent protein (GFP) dual-expressing MM.1S cells were injected into mice via the tail vein four weeks after irradiation (**Fig. 4I, Fig. S9G**). Vehicle, pomalidomide, or AU-24118 were then orally administered. The luciferase signal showed a substantial reduction over time and at endpoint, indicative of diminished tumor proliferation (**Fig. 4J, Fig. S9H**). A notable extension in the overall survival of mice treated with AU-24118 was observed (**Fig. 4K**), and TUNEL staining was significantly increased following AU-24118 treatment (**Fig. 4L**). IHC confirmed loss of SMARCA4 and c-MYC exclusively in AU-24118-treated tumors in both the MM1.S disseminated and subcutaneous models (**Fig. S9I-J**).

Histopathological evaluation of orthotopic xenografts to assess the efficacy of the mSWI/SNF ATPase degrader was undertaken. Pathological assessment revealed that in comparison to the vehicle where sheets of plasma cells were noted, there was an absence of any perceptible plasma cells in the AU-24118-treated group. Also, in AU-24118-treated tumors, we identified remnant hematopoietic cells intermixed (not seen in vehicle tumor tissues) with a fair number of red blood cell (RBC)-filled sinusoidal areas. The presence of areas filled with RBCs in the sinusoids in the marrow tissue, which appear to be areas of drug-mediated tumor regression, along with the presence of hematopoietic cells, provides additional direct (*in situ*) biological evidence of the better efficacy of our degrader (**Fig. 4M**). This was in turn validated molecularly on CD38 IHC wherein comparison to diffuse strong membranous positivity of CD38 in all the marrow cells of the vehicle tumor tissue, we saw near total absence of CD38 in any remnant cells in the AU-24118-treated orthotopic xenografts. This points towards a significant and complete abatement of tumor cells upon AU-24118 treatment. Additionally, a standard of care therapeutic (pomalidomide) showed some depletion of plasma cells but not a degree of depletion as seen in the AU-24118-treated group at both morphological and molecular levels (**Fig. 4M**).

## Discussion

Transcription factors are frequently dysregulated in the pathogenesis of human cancer, representing a major class of cancer cell dependencies. Targeting these factors can significantly impact the treatment of specific malignancies, as exemplified by the clinical success of agents targeting the androgen receptor (AR) in prostate cancer and estrogen receptor (ER) in breast cancer^36^. Conventional small-molecule drugs exert their effects by binding to defined pockets on target protein surfaces, such as the ligand binding domains of AR and ER. However, many TFs lack structurally ordered ligand binding pockets, presenting significant challenges in therapeutically targeting their actions. As an alternative strategy, targeting of TF coregulators has emerged as a promising approach to block their functions in cancer^37^. We previously found that inhibiting the mSWI/SNF chromatin remodeling complex disrupts oncogenic signaling of key TFs (AR, FOXA1, ERG, and MYC) in castration-resistant prostate cancer (CRPC)^23^. Here, we identify the mSWI/SNF complex as a therapeutic vulnerability in other TF-driven malignancies, namely POU2F3-driven SCLC and POU2AF1-dependent B cell malignancies. Importantly, we show that an orally bioavailable mSWI/SNF ATPase degrader, AU-24118, has anti-tumor activity in multiple preclinical models of both SCLC-P and MM with no signs of toxicity.

Our findings identify a pronounced dependency of SCLC-P cells, but not cells of the other molecular subtypes, on the mSWI/SNF complex, highlighting a critical epigenetic regulatory axis for POU2F3 signaling. The data from our CRISPR screen, contrasting the dependency of SCLC-A cells and SCLC-P cells on the mSWI/SNF complex, could be partially explained by the association between the POU2F3-POU2AF2/3 complex with the mSWI/SNF complex. The findings also invite further inquiry into the regulatory mechanisms underlying ASCL1’s transcriptional activity in SCLC-A cells as ASCL1 may rely on alternative mechanisms to modulate chromatin accessibility in SCLC-A cells. Moreover, the therapeutic efficacy of mSWI/SNF ATPase degraders in SCLC-P suggests broader implications whereby other POU2F3-expressing small cell carcinomas may respond to this targeted therapy. Notably, SCLC shares transcriptional drivers with neuroendocrine prostate cancer (NEPC)^38^, and the mSWI/SNF complex has been suggested to be involved in NEPC^39^. Recently, multiple single-cell analyses have identified a subpopulation of NEPC cells with high expression of POU2F3 and its downstream target ASCL2 in both prostate cancer patients and genetically engineered mouse models (GEMMs)^40–43^. As androgen deprivation therapy (ADT) continues to be a standard treatment for prostate cancer, the emergence of NEPC post-ADT underscores the need to explore mSWI/SNF targeting therapies in POU2F3-expressing NEPC.

In addition to SCLC-P cells, this study reveals that mSWI/SNF ATPase degraders have potent therapeutic activity against a subset of MM and DLBCL cells that are dependent on the POU2AF1 coactivator. Multi-omics analysis unveils a novel role of POU2AF1 as a coactivator for IRF4, which also has been implicated by others^44^. mSWI/SNF ATPase degrader treatment effectively compacts chromatin in MM cells, dislodging IRF4 and POU2AF1 from DNA and decreasing oncogenic IRF4 signaling. Given the critical function of IRF4 in B cell malignancies and the lack of FDA-approved strategies directly targeting IRF4, these findings have significant impact by providing an alternative avenue of therapeutic intervention through inhibiting mSWI/SNF and the actions of the POU2AF1 coactivator.

The embryonic lethality observed upon genetic knockout of the ATPase subunit of the mSWI/SNF complex necessitates a thorough examination of the toxicity profile associated with ATPase subunit degradation *in vivo*^45,46^. Our *in vivo* assessments with the first-in-class orally bioavailable SMARCA2/4 PROTAC degrader, AU-24118, demonstrated a favorable tolerability profile alongside significant anti-tumor efficacy in multiple SCLC-P and MM preclinical models. Moreover, in the *in vivo* models of SCLC-P, AU-24118 treatment did not affect tuft cells in normal tissues. Effective regenerative processes were also observed in disseminated orthotopic xenograft models of MM, addressing concerns regarding potential adverse effects on normal cellular processes. Similar observations were made by Papillon et al., where hematopoietic stem cells (HSC) isolated from BRM014 (SMARCA2/4 inhibitor)^47^-treated mice retained their functionality, suggesting transient loss of mSWI/SNF function does not permanently suppress HSC function^25^. Recent studies delineating the role of the mSWI/SNF complex in memory T cell fate suggest that modulating mSWI/SNF activity early in T cell differentiation can enhance cancer immunotherapy outcomes^48,49^, thereby warranting future studies to evaluate the anti-tumor efficacy and safety of mSWI/SNF-targeting strategies in syngeneic tumor models in immunocompetent mice.

Collectively, this study identifies the mSWI/SNF chromatin remodeling complex as a novel vulnerability in POU2F3-dependent SCLC and POU2AF1-dependent MM (**Fig. 5**). Combined with our previous findings in CRPC^23^, this suggests that mSWI/SNF-targeted therapeutics may have efficacy across a range of cancer types driven by select TFs. These data position mSWI/SNF ATPase PROTAC degraders for further optimization, development, and testing in clinical settings.

**Figure 5.**
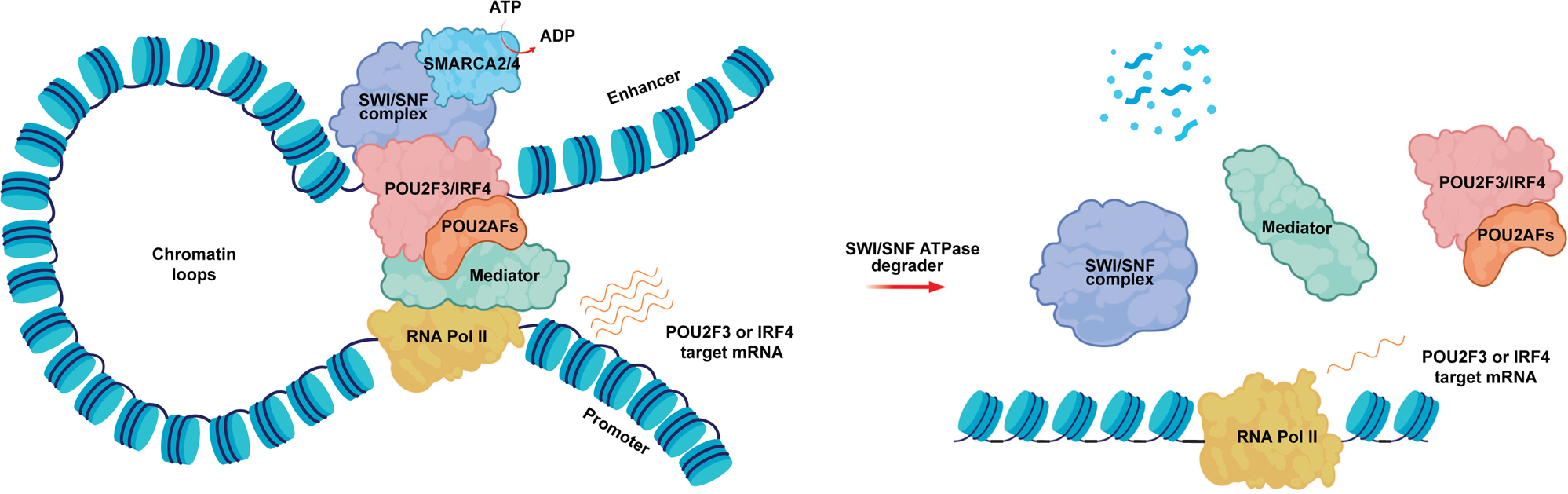
Schematic showing the mechanism of action of mSWI/SNF ATPase degraders in POU2F3-dependent SCLC and POU2AF1-dependent B cell malignancies. The mSWI/SNF complex remodels nucleosomes at enhancer sites, allowing physical access to transcription factors (TFs) -POU2F3 in SCLC-P and IRF4 in multiple myeloma. Activity of the TFs is regulated by requisite coactivators (POU2AFs). This leads to recruitment of the transcriptional machinery at promoters, driving transcription of POU2F3 or IRF target genes. SMARCA2/4 ATPases are degraded with PROTAC treatment, resulting in loss of mSWI/SNF complex activity and eviction of TFs/coactivators and transcriptional machinery. This leads to diminished transcription of POU2F3 and IRF4 target genes and decreased oncogenic signaling from these pathways.

## STAR Methods

### Cell lines, antibodies, and compounds

All cell lines were originally obtained from ATCC, DSMZ, ECACC or internal stock. All cell lines were genotyped to confirm their identity at the University of Michigan Sequencing Core and tested biweekly for Mycoplasma contamination. H526, H1048, H211, COR-311, MM1S, and Karpas-25 were grown in Gibco RPMI-1640 + 10% fetal bovine serum (FBS) (ThermoFisher Scientific). H929 was grown in Gibco RPMI-1640 + 10% FBS + 0.05mM 2-mercaptoethanol. Sources of all antibodies are described in **Supplementary Table 2**. AU-15330 and AU-24118 were designed and synthesized by Aurigene Oncology. Lenalidomide and pomalidomide were purchased from Selleck Chemicals. Carfilzomib was purchased from the Michigan Medicine pharmacy.

### Paralog gene identification and functional domain mapping

Paralog pairs within the human genome were identified using BlastP. Matches of isoforms originating from the same gene were removed. Each individual gene’s top paralog identified (E-value < 0.01) that shared the same functional domain of interest was included in the Paralog library. In addition, each paralog pair was included for genes with multiple high-scoring paralogs (E-value < 10-100). Functional domains were mapped using reverse spi blast (rps-Blast) and the conserved domain database (CDD)^50^.

### Selection of sgRNAs and controls

Domain annotation and sgRNA cutting codon were compared, and sgRNAs cutting in functional domain regions were included in the sgRNA selection pool. sgRNAs with off-targets in paralog genes were removed from the selection pool. sgRNAs were picked based on their off-target score (calculated based on the number of off-target locations in the human genome and number of miss-matches). For each gene, 3-4 selective domain-focused sgRNA were picked. In cases in which selective domain-focused targeting sgRNA were not available, sgRNAs targeting the upstream coding region of the gene were selected. For each given paralog pair (A-B), 3-4 sgRNA for paralog A were combined with 3-4 sgRNAs for paralog B, resulting in 9-16 combinations. To evaluate single-gene knockout effects of each gene, each of the paralog’s sgRNA was also combined with each one targeting- and one non-targeting-negative control. A set of known essential genes as positive controls (dgRNA n=28) and a set of non-targeting (dgRNA n=100) as well as non-coding region targeting negative controls (dgRNA n=54) were generated. To construct cell line-specific negative controls (non-synergistic pairs), we selected genes that were not expressed in a cell line according to the RNA sequencing (RNA-seq) data (log2(TPM + 1) < 0.1) from the CCLE.

### CRISPR screening library generation

The paralog co-targeting CRISPR library was generated to use SpCas9, a system recently published^51^.Oligonucleotide pools, targeting 4,341 single genes and 4,387 paralogs using 137,950 double guide RNAs, were synthesized (Twist Bioscience) and cloned into LRG3.0, a lentiviral vector with human U6 and bovine U6 promoters expressing the two sgRNAs in inverse orientation. Cas9 stable cell lines were transduced with Cas9 vector (Addgene: 108100). Cell lines were transduced with the paralog co-targeting CRISPR library virus to achieve a representation of 1,000 cells per sgRNA at a low multiplicity of infection (around 0.3). SCLC cell lines were transduced while spun for 45 min at 600g. On day 6 after transduction, cells were selected using blasticidin, split, and replated to maintain representation. An initial sample was taken using the remainder. Once 10 cell doublings were reached, cells were pelleted by centrifugation and frozen, or genomic DNA was extracted directly.

### Genomic DNA extraction

Cells were resuspended in resuspension buffer (10mM Tris-HCl pH=8.0, 150mM NaCl, 10mM EDTA) with the addition of proteinase K (0.02mg/mL) and SDS (final concentration 0.1%). Lysate was incubated at 56°C for 48h. Genomic DNA was extracted using two rounds of TRIS-saturated phenol (Thermo Fisher Scientific) extraction.

### dgRNA PCR for Illumina sequencing

For PCR from genomic DNA, 1µg of genomic DNA was used for each reaction. In round 1, PCR with 11 cycles was used. DNA was purified using a gel extraction kit (QIAGENE) according to the manufacturer’s instructions. Product DNA was barcoded by amplification in a second round PCR using stacked P5/P7 primers. PCR products were again purified and sequenced on NextSeq with the paired-end 75 base pair (bp) reads protocol (Illumina). Reads were counted by mapping the pairs of 19–20 nt sgRNAs to the reference sgRNA list containing combinations present in the library. 16 pseudo counts were added prior to downstream analysis. The resulting matrix of read counts was used to calculate log2 fold changes.

### Calculation of paralog CRISPR screening Log_2_ fold changes and synergy scores

Synergy scores were calculated using the GEMINI R package^52^ (**Supplementary Table 1**). Briefly, GEMINI calculates the LFC of the sgRNA pair abundance between the initial- and the 10-doubling time endpoint. GEMINI has been used to compute the synergy score by comparing the LFC of each gene pair to the most lethal individual gene of the pair. GEMINI uses non-synergistic pairs to calculate the FDR and p-value in each cell line, as described previously^52^. Beta scores for single and double knockouts were calculated using MAGeCK^26,52^ and compared between 3 SCLC-A and 3 SCLC-P cell lines. Gene-level beta scores for synergistic double gene knockouts (synergy score > 1) (n=968) and single knockouts were plotted.

### Cell viability assay

Cells were plated onto 96-well plates in their respective culture medium and incubated at 37 °C in an atmosphere of 5% CO_2_. After overnight incubation, a serial dilution of compounds was prepared and added to the plate. The cells were further incubated for 5 days, and the CellTiter-Glo assay (Promega) was then performed according to the manufacturer’s instruction to determine cell proliferation. The luminescence signal from each well was acquired using the Infinite M1000 Pro plate reader (Tecan), and the data were analyzed using GraphPad Prism software (GraphPad Software).

### Western blot

Western blot was performed as previous described^23^. In brief, cell lysates were prepared in RIPA buffer (ThermoFisher Scientific) supplemented with protease inhibitor cocktail tablets (Sigma-Aldrich). Total protein concentration was measured by Pierce BCA Protein Assay Kit (ThermoFisher Scientific), and an equal amount of protein was loaded in NuPAGE 3 to 8% Tris-Acetate Protein Gel (ThermoFisher Scientific) or NuPAGE 4 to 12% Bis-Tris Protein Gel (ThermoFisher Scientific) and blotted with primary antibodies. Following incubation with HRP-conjugated secondary antibodies, membranes were imaged on an Odyssey CLx Imager (LiCOR Biosciences). For all immunoblots, uncropped and unprocessed images are provided in **Fig. S10**.

### RNA isolation and quantitative real-time PCR

Total RNA was isolated from cells using the Direct-zol kit (Zymo), and cDNA was synthesized using Maxima First Strand cDNA Synthesis Kit for PCR with reverse transcription (RT–PCR) (Thermo Fisher Scientific). Quantitative real-time PCR (qPCR) was performed in triplicate using standard SYBR green reagents and protocols on a QuantStudio 7 Real-Time PCR system (Applied Biosystems). The target mRNA expression was quantified using the ΔΔCt method and normalized to *ACTB* expression. Primer sequences are listed in **Supplementary Table 2**.

### ATAC-seq and analysis

ATAC-seq was performed as previously described^53^. In brief, cells treated with AU-15330 were washed in cold PBS and resuspended in RSB buffer with NP-40, Tween-20, protease inhibitor and digitonin cytoplasmic lysis buffer (CER-I from the NE-PER kit, Thermo Fisher Scientific). This single-cell suspension was incubated on ice for 5 min. The lysing process was terminated by the addition of double volume RSB buffer with Tween-20. The lysate was centrifuged at 1,300g for 5 min at 4 °C. Nuclei were resuspended in 50 μl of 1× TD buffer, then incubated with 0.5–3.5 μl Tn5 enzyme for 30 min at 37 °C (Nextera DNA Library Preparation Kit; cat. no. FC-121-1031). Samples were immediately purified by Qiagen minElute column and PCR-amplified with the NEB Next High-Fidelity 2X PCR Master Mix (cat. no. M0541L) following the original protocol. qPCR was used to determine the optimal PCR cycles to prevent over-amplification. The amplified library was further purified by Qiagen minElute column and SPRI beads (Beckman Coulter, cat. no. A63881). ATAC-seq libraries were sequenced on the Illumina HiSeq 2500 or NovaSeq. fastq files were trimmed using Trimmomatic (version 0.39) and then uniquely aligned to the GRCh38/hg38 human genome assembly using bwa mem (version 0.7.17-r1198-dirty) and converted to binary files using SAMtools (version 1.9)^54–56^. Reads mapped to mitochondrial or duplicated reads were removed by SAMtools and PICARD MarkDuplicates (version 2.26.0-1-gbaf4d27-SNAPSHOT), respectively. Filtered alignment files from replicates were merged for downstream analysis. MACS2 (2.1.1.20160309) was used to call ATAC-seq peaks [IDS: Macs]. UCSC’s tool wigtoBigwig was used for conversion to bigwig formats [UCSC 20639541]. All de novo and known motif enrichment analyses were performed using the HOMER (version v4.11.1) suite of algorithms^57^. De novo motif discovery and enrichment analysis of known motifs were performed with findMotifsGenome.pl (–size given).

### RNA-seq and analysis

RNA-seq libraries were prepared using 200–1,000 ng of total RNA. PolyA+ RNA isolation, cDNA synthesis, end-repair, A-base addition, and ligation of the Illumina indexed adapters were performed according to the TruSeq RNA protocol (Illumina). Libraries were size selected for 250– 300 bp cDNA fragments on a 3% Nusieve 3:1 (Lonza) gel, recovered using QIAEX II reagents (QIAGEN), and PCR amplified using Phusion DNA polymerase (New England Biolabs). Library quality was measured on an Agilent 2100 Bioanalyzer for product size and concentration. Paired-end libraries were sequenced with the Illumina HiSeq 2500, (2 × 100 nucleotide read length) with sequence coverage to 15–20M paired reads. Libraries passing quality control were trimmed of sequencing adaptors and aligned to the human reference genome, GRCh38. Samples were demultiplexed into paired-end reads using Illumina’s bcl2fastq conversion software v2.20. The reference genome was indexed using bwa (version 0.7.17-r1198-dirty), and reads were pseudoaligned onto the GRCh38/hg38 human reference genome using Kallisto’s quant command^54,58^. EdgeR (version 3.39.6) was used to compute differential gene expression using raw read-counts as input^59^. Limma-Voom (limma_3.53.10) was then used to perform differential expression analysis^60^. Heatmaps were generated using the ComplexHeatmap package in R. These gene signatures were used to perform a fast pre-ranked GSEA using fgsea bioconductor package in R (version fgsea_1.24.0). We used the function fgsea to estimate the net enrichment score and p-value of each pathway, and the plotEnrichment function was used to plot enrichment for the pathways of interest.

### POU2F3/AF2/AF3-dTAG-HA system expressing SCLC cells

For HA-dTAG-POU2F3 or POU2AF2/3-dTAG-HA system, the FKBP23F36V-2xHA was PCR amplified from the pCRIS-PITCHv2-Puro-dTAG vector (Addgene: 91703) and introduced into sgRNA-resistant POU2F3_LentiV_neo or the POU2AF2/3_LentiV_neo vector for functional validation with competition-based cell proliferation assay. NCI-H1048/NCI-H526 that stably expressed Cas9 were infected either with HA_dTAG_POU2F3_LentiV_neo or POU2AF2/3_dTAG_HA_LentiV_neo or empty_vector_lentiV_neo construct followed by neomycin selection to establish stable cell lines. The cells were then lentivirally delivered with indicated sgRNAs co-expressed with a GFP reporter. The percentage of GFP+ cells correspond to the sgRNA representation within the population. GFP measurements in human cell lines were taken on day 4 post-infection and every four days with Guava Easycyte HT instrument (Millipore). The fold change in GFP+ population (normalized to day 4) was used for analysis. HA_dTAG_POU2F3 or POU2AF2/3_dTAG_HA, which is resistant to its own sgRNA, were cloned into the LRGB2.1T vector that either contains sgRNA against endogenous POU2F3 or POU2AF2/3 into NCIH211/NCIH526/NCIH1048 that stably express Cas9.

### ChIP–seq and data analysis

Chromatin immunoprecipitation experiments were carried out using the ideal ChIP-seq kit for TFs (Diagenode) as per the manufacturer’s protocol. Chromatin from 2 × 10^6^ cells was used for each ChIP reaction with 4μg of the target protein antibody. In brief, cells were trypsinized and washed twice with 1× PBS, followed by cross-linking for 10 min in 1% formaldehyde solution. Crosslinking was terminated by the addition of 1/10 volume 1.25 M glycine for 5 min at room temperature followed by cell lysis and sonication (Bioruptor, Diagenode), resulting in an average chromatin fragment size of 200 bp. Fragmented chromatin was then used for immunoprecipitation using various antibodies, with overnight incubation at 4 °C. ChIP DNA was de-crosslinked and purified using the standard protocol. Purified DNA was then prepared for sequencing as per the manufacturer’s instructions (Illumina). ChIP samples (1–10 ng) were converted to blunt-ended fragments using T4 DNA polymerase, Escherichia coli DNA polymerase I large fragment (Klenow polymerase), and T4 polynucleotide kinase (New England BioLabs (NEB)). A single adenine base was added to fragment ends by Klenow fragment (3′ to 5′ exo minus; NEB), followed by ligation of Illumina adaptors (Quick ligase, NEB). The adaptor-ligated DNA fragments were enriched by PCR using the Illumina Barcode primers and Phusion DNA polymerase (NEB). PCR products were size-selected using 3% NuSieve agarose gels (Lonza) followed by gel extraction using QIAEX II reagents (Qiagen). Libraries were quantified and quality checked using the Bioanalyzer 2100 (Agilent) and sequenced on the Illumina HiSeq 2500 Sequencer (125-nucleotide read length). Paired-end, 125 bp reads were trimmed and aligned to the human reference genome (GRC h38/hg38) with the Burrows-Wheeler Aligner (BWA; version 0.7.17-r1198-dirty) The SAM file obtained after alignment was converted into BAM format using SAMTools (version 1.9)^56^. Picard MarkDuplicates command and samtools were used to filter aligned output. MACS2 (version 2.1.1.20160309) callpeak was used for performing peak calling with the following option: ‘macs2 callpeak–call-summits–verbose 3 -g hs -f BAM -n OUT–qvalue 0.05^61^. Blacklisted regions of the genome were removed using bedtools. UCSC’s tool wigtoBigwig was used for conversion to bigwig formats. ChIP peak profile plots and read-density heat maps were generated using deepTools, and cistrome overlap analyses were carried out using the ChIPpeakAnno (version 3.0.0) or ChIPseeker (version 1.29.1) packages in R (version 3.6.0)^62–64^.

### FPLC

NCI-H526/COR-L311 nuclear extracts were obtained using NE-PER nuclear extraction kit (Thermo Scientific) and dialyzed against FPLC buffer (20 mM Tris-HCl, 0.2 mM EDTA, 5mM MgCl2, 0.1 M KCl, 10% (v/v) glycerol, 0.5 mM DTT, 1 mM benzamidine, 0.2 m MPMSF, pH7.9). 5mg of nuclear protein was concentrated in 500 μl using a Microcon centrifugal filter (Millipore) and then applied to a Superose 6 size exclusion column (10/300 GL GE Healthcare) pre-calibrated using the Gel Filtration HMW Calibration Kit (GE Healthcare). 500 μl elute was collected for each fraction at a flow rate of 0.5ml/min, and eluted fractions were subjected to SDS-PAGE and western blotting.

### RIME and data analysis

RIME experiments were carried out as previously described^65^. In brief, 40 × 10^6^ cells were used for each RIME reaction with 20 μg of the target protein antibody. Cells were harvested followed by cross-linking for 8 min in 1% formaldehyde solution. Crosslinking was terminated by adding glycine to a final concentration of 0.1M for 5 min at room temperature. Cells were washed with 1x PBS and pelleted by centrifugation at 2000g for 3 min at 4°C for 4 times total. Cell pellets were added to the nuclear extraction buffer LB1, LB2, and LB3 separately. Lysates were sonicated (Bioruptor, Diagenode) to result in an average chromatin fragment size of 200-600 bp. Fragmented nuclear lysates were then used for immunoprecipitation using various antibodies, with overnight incubation at 4 °C. All antibodies were preincubated with beads for 1 hour at room temperature. Total protein per replicate was labeled with TMT isobaric Label Reagent (Thermo Fisher Scientific) according to the manufacturer’s protocol and subjected to liquid chromatography−mass spectrometry (LC−MS)/MS analysis.

### AU-24118, pomalidomide, and carfilzomib formula for *in vivo* studies

AU-24118 was added in PEG200 and then sonicated and vortexed until completely dissolved. Five volumes of 10% D-α-Tocopherol polyethylene glycol 1000 succinate was next added, and the solution was vortexed until homogeneous. Four volumes of 1% Tween 80 was then added, and the solution was vortexed until homogeneous. AU-24118 was freshly prepared right before administration to mice. Pomalidomide was dissolved in DMSO and then added in 30% PEG400 + 2%Tween 80 + 68% ddH_2_O. AU-24118 and pomalidomide were delivered to mice by oral gavage. Carfilzomib was diluted in sterile water based on the company’s instructions (Kyprolis).

### Human tumor xenograft models

Six-week-old male CB17 severe combined immunodeficiency (SCID) mice were procured from the University of Michigan breeding colony. Subcutaneous tumors were established at both sides of the dorsal flank of mice. Tumors were measured at least biweekly using digital calipers following the formula (π/6) (L × W^2^), where L is length and W is width of the tumor. The disseminated model was measured by signal intensity of luminescence by PerkinElmer’s IVIS Spectrum from the University of Michigan imaging core. At the end of the studies, mice were killed and tumors extracted and weighed. The University of Michigan Institutional Animal Care and Use Committee (IACUC) approved all *in vivo* studies. For the H526, H1048, and H69 models, 5 × 10^6^ tumor cells were injected subcutaneously into the dorsal flank on both sides of the mice in a serum-free medium with 50% Matrigel (BD Biosciences). Once tumors reached a palpable stage (∼100 mm^3^), mice were randomized and treated with either 15 mg kg^−1^ AU-24118 or vehicle by oral gavage 3 days per week for 3 - 4 weeks. For the H929 and MM1S models, 5 × 10^6^ cells were injected subcutaneously into the dorsal flank on both sides of the mice in a serum-free medium with 50% Matrigel (BD Biosciences). Once tumors reached a palpable stage (∼100 mm^3^), mice were randomized and treated with the following as indicated in the figures: 15 mg kg^−1^ AU-24118 by oral gavage 3 days per week, 10 mg kg^−1^ pomalidomide by oral gavage 5 days per week, 5 mg kg^−1^ carfilzomib by intravenous administration for two consecutive days and 5 days rest, or vehicle for 3-4 weeks. For the Karpas-25 tumor model, 3 × 10^6^ cells were injected subcutaneously into the dorsal flank on both sides of the mice in a serum-free medium with 50% Matrigel (BD Biosciences). Once tumors reached a palpable stage (∼100 mm^3^), mice were randomized and treated with either 15 mg kg^−1^ AU-24118 by oral gavage 3 days per week, 5 mg kg^−1^ carfilzomib by intravenous injection for two consecutive days injection and 5 days rest, or vehicle for 3-4 weeks. For the MM1S disseminated model, 1 × 10^7^ GFP/luc MM1.S cells were injected intravenously from the tail vein of the mice in a PBS medium after 24 hours 250 cGy r-irradiation using the Small Animal Radiation Research Platform (SARRP). The mice were then treated with 1 mg/ml neomycin water bottle for 3 weeks in case of infection due to irradiation. Once the signal of luminescence reached a measurable stage (∼1 × 10^6^), mice were randomized and treated with either 15 mg kg^−1^ AU-24118 by oral gavage 3 days per week, pomalidomide by oral gavage 5 days per week, or vehicle until the mice reached the endpoint based on protocol. Following the IACUC guidelines, in all treatment arms, the maximal tumor size did not exceed the 2.0 cm limit in any dimension, and animals with xenografts reaching that size were duly euthanized.

### GFP/Luc MM1.S cell line

MM1.S cells were transduced with GFP luciferase lentivirus (purchased from the vector core of University of Michigan) through spinfection (45 minutes at 600g). Two days after the viral transduction, the GFP-positive cells were sorted with a cell sorter (SONY SH800S).

### Histopathological analysis of organs harvested for drug toxicity

For the present study, organs (liver, spleen, kidney, small intestine, and lung) were harvested and fixed in 10% neutral buffered formalin followed by embedding in paraffin to make tissue blocks. These blocks were sectioned at 4 µm and stained with Harris haematoxylin and alcoholic eosin-Y stain (both reagents from Leica Surgipath), and staining was performed on a Leica autostainer-XL (automatic) platform. The stained sections were evaluated by two different pathologists using a brightfield microscope in a blinded fashion between the control and treatment groups for general tissue morphology and coherence of architecture. A detailed comprehensive analysis of the changes noted at the cellular and subcellular level were performed as described below for each specific tissue. Evaluation of liver: Liver tissue sections were evaluated for normal architecture, and regional analysis for all three zones was performed for inflammation, necrosis, and fibrosis. Evaluation of spleen: Splenic tissue sections were evaluated for the organization of hematogenous red and lymphoid white pulp regions including necrosis and fibrotic changes, if any. Evaluation of kidney: Kidney tissue sections were examined for changes noted, if any, in all four renal functional components, namely glomeruli, interstitium, tubules, and vessels. Evaluation of small intestine: Small intestine tissue sections were examined for mucosal changes such as villous blunting, villous: crypt ratio, and evaluated for inflammatory changes including intraepithelial lymphocytes, extent (mucosal, submucosal, serosal), and type of inflammatory infiltrate including tissue modulatory effect. Evaluation of lung: Lung tissue sections were thoroughly examined to identify the presence of regenerative/degenerative atypia in the alveolar and bronchiolar epithelium, hyperplasia of type II pneumocytes, and interstitial pneumonia. The presence of extensive alveolar damage, organized pneumonia (also known as bronchiolitis obliterans organizing pneumonia or BOOP), and alveolar hemorrhage and histology suggesting usual interstitial pneumonitis (UIP) was also investigated. A mild and within normal range proliferation of type II pneumocytes (devoid of other associated inflammatory and other associative findings) was considered within unremarkable histology.

### Immunohistochemistry

Immunohistochemistry (IHC) was performed on 4-micron formalin-fixed, paraffin-embedded (FFPE) tissue sections using POU2F3, BRG1 (a surrogate marker for SMARCA4), CD38, and DCLK1. IHC was carried out on the Ventana ULTRA automated slide staining system using the Omni View Universal DAB detection kit. The antibody and critical reagent details have been provided in **Supplementary Table 2.** Either the presence or absence of BRG1 and POU2F3 nuclear staining and DCLK1 and CD38 cytoplasmic/membranous staining were recorded by the study pathologists. To provide a semi-quantitative score per biomarker, a product score was rendered wherever needed. The IHC product score calculated out of 300 was derived by multiplying the percentage of positive tumor cells (PP) for each staining intensity (I) and adding the values in each tumor using the formula “IHC Score = (PP * 0 + PP * 1 + PP * 2 + PP * 3)” as previously described^66^.

### Specialized IHC tissue modulatory score for normal organs

To rule out modulatory effects on the molecular levels as predicted by unremarkable morphology on histopathological assessment of the normal organs, a specialized histology score was devised to fit the individual organ systems. For the intestine, the number of DCLK1-positive cells/ 500 intestinal enterocytes (predominantly villi of small intestine) were counted; for lung parenchyma, the number of DCLK1-positive cells/5 high power fields were counted.

### TUNEL assay

Apoptosis was examined using Terminal dUTP Nick End Labeling (TUNEL) performed with an In Situ Cell Death Detection Kit (TMR Red #12156792910; Roche Applied Science) following the manufacturer’s instructions. Briefly, fixed sections were permeabilized with Triton X-100, followed by a PBS wash. The labeling reaction was performed at 37 °C for 60 min by addition of a reaction buffer containing enzymes. Images were acquired on a Zeiss Axiolmager M1 microscope.

## Data and code availability

ATAC-seq, ChIP-seq, and RNA-seq data have been deposited at the Gene Expression Omnibus (GEO), accession number GSE247951. Any additional information required to reanalyze the data reported in this work is available from the lead contact upon request.

## Acknowledgements

We thank Lisa McMurry, Rui Wang, Amanda Miller, Christine Caldwell-Smith, Xia Jiang, Sarah Yee, Yunhui Cheng, Rupam Bhattacharyya, and Shuqin Li from the Michigan Center for Translational Pathology at the University of Michigan as well as Venkatesha Basrur and the Rogel Cancer Center Proteomics Shared Resource. This work was funded primarily by the Trailsend Foundation with additional support from the Prostate Cancer Foundation (PCF), National Cancer Institute (NCI) Prostate Specialized Programs of Research Excellence (SPORE) Grant P50-CA186786, and an NCI Outstanding Investigator Award R35-CA231996 (A.M.C.). L.X. is a recipient of the 2022 PCF Young Investigator Award, and he is also supported by a Department of Defense Prostate Cancer Research Program Idea Development Award (W81XWH-21-1-0500) and the Michigan SPORE Career Enhancement Program. A.M.C. is a Howard Hughes Medical Institute Investigator, A. Alfred Taubman Scholar, and American Cancer Society Professor.

## Author contributions

L.X., A.M.C., C.R.V., and T.H. designed and conceived the project. T.H. and L.X. performed all *in vitro* and functional genomic experiments with assistance from Y.Q., A.P., S.E., S.H., X.W., N.H.K., and H.Z. O.K., X.S.W, and C.R.V. performed the CRISPR screening and generated the dTAG cell line. T.H. performed all animal efficacy studies with help from Y.Q., Y.Z., and J.C.T. T.H., L.X., and E.Y. carried out all bioinformatic analyses with assistance from A.P. R. Mannan, S.M., and R. Mehra carried out all histopathological evaluations of drug toxicity as well as quantified all histology-based data and immunohistochemistry. F.S. and X.C. generated next-generation sequencing libraries and performed the sequencing. C.A., S.S., and M.R. contributed to the discovery of AU-15330 and AU-24118 compounds. T.H., L.X., S.J.M., and A.M.C. wrote the manuscript and organized the final figures. All authors read, commented, and participated in manuscript review and editing.

## Declaration of Interests

A.M.C. serves on the clinical advisory board of Aurigene Discovery Technologies. C.A., S.S., and M.R. are employees of Aurigene Discovery Technologies. C.R.V. has received consulting fees from Flare Therapeutics, Roivant Sciences, and C4 Therapeutics; he has served on the advisory boards of KSQ Therapeutics, Syros Pharmaceuticals, and Treeline Biosciences. C.R.V. has also received research funding from Boehringer-Ingelheim and Treeline Biosciences and owns stock in Treeline Biosciences. Aurigene has filed patent applications on AU-15330 and AU-24118. The other authors declare no competing interests.

## Notes

### Competing Interest Statement

A.M.C. serves on the clinical advisory board of Aurigene Discovery Technologies. A.C.S., S.S., and M.R. are employees of Aurigene Discovery Technologies. C.R.V. has received consulting fees from Flare Therapeutics, Roivant Sciences, and C4 Therapeutics; he has served on the advisory boards of KSQ Therapeutics, Syros Pharmaceuticals, and Treeline Biosciences. C.R.V. has also received research funding from Boehringer-Ingelheim and Treeline Biosciences and owns stock in Treeline Biosciences. Aurigene has filed patent applications on AU-15330 and AU-24118. The other authors declare no competing interests.

### Summary of Updates

The revised manuscript contains a correction to Figure S5C and a correction to an author's name.

