## Supplemental Figures 1 - 10 for "Targeting the mSWI/SNF Complex in POU2F-POU2AF Transcription Factor-Driven Malignancies"

### Figure S1

A

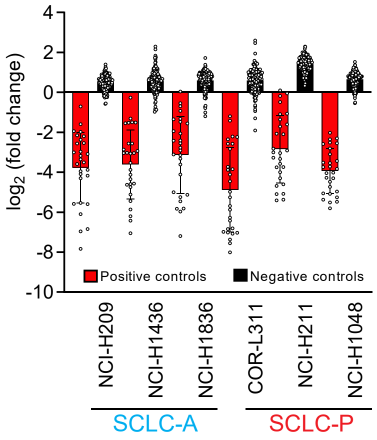


C

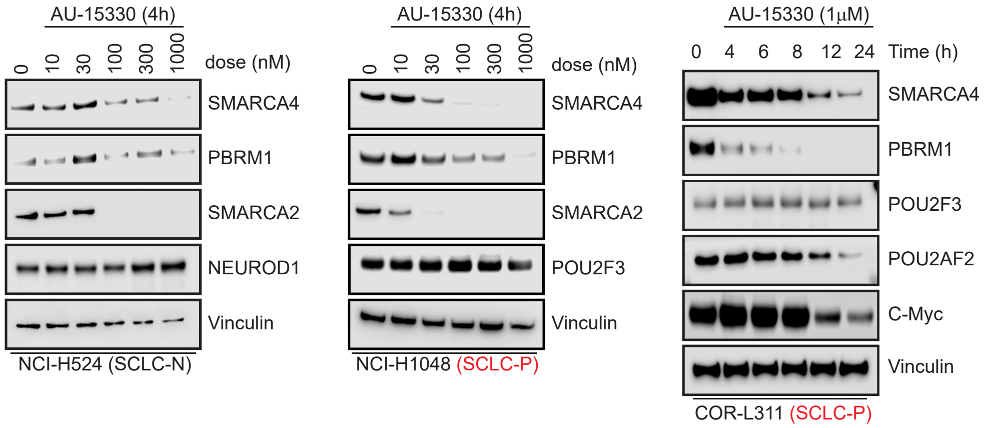


B

| sgRNA combinations | sgRNA number | SCLC-P | | | SCLC-A | | | Beta score |
|---|---|---|---|---|---|---|---|---|
| | | COR-L311 | NCI-H1048 | NCI-H211 | NCI-H1436 | NCI-H1836 | NCI-H209 | |
| sgSMARCA2 (Bromodomain) + sgControl | 8 | -1.19 | 0.07 | -1.09 | -0.03 | -0.18 | -0.4 | -0.54 |
| sgSMARCA4 (Bromodomain) + sgControl | 8 | -1.83 | -0.73 | -2.12 | -0.2 | -0.26 | -0.33 | -1.29 |
| sgSMARCA2 (Bromodomain) + sgSMARCA4 (Bromodomain) | 16 | -2.13 | -1.00 | -2.28 | -0.37 | -0.7 | -0.66 | -1.22 |
| sgSMARCA2 (RRM domain) + sgControl | 8 | -0.39 | -0.08 | -0.73 | 0.32 | -0.09 | -0.05 | -0.46 |
| sgSMARCA4 (RRM domain) + sgControl | 8 | -2.76 | -1.39 | -2.14 | -0.88 | -0.66 | -0.71 | -1.35 |
| sgSMARCA2 (RRM domain) + sgSMARCA4 (RRM domain) | 16 | -2.14 | -1.19 | -2.31 | -1.05 | -0.92 | -1.24 | -0.81 |
| sgSMARCA2 (Bromodomain) + sgPBRM1 (Bromodomain) | 16 | -1.62 | -0.24 | -1.39 | -1.11 | -0.73 | -0.77 | -0.21 |
| sgBRD9 (Bromodomain) + sgControl | 8 | -2.64 | -0.49 | -1.89 | -0.36 | -0.51 | -1.09 | -1.02 |
| sgBRD9 (Bromodomain) + sgBRD7 (Bromodomain) | 12 | -3.32 | -1.15 | -2.17 | -0.59 | -1.03 | -1.54 | -1.16 |

D

| SCLC cell line | Subtype | AU-15330 (nM) |
|---|---|---|
| NCI-H526 | SCLC-P | 1.8 |
| COR-L311 | SCLC-P | 8.86 |
| NCI-H1048 | SCLC-P | 11.45 |
| NCI-H211 | SCLC-P | 21.91 |
| NCI-H69 | SCLC-A | 4406 |
| NCI-H1963 | SCLC-A | 5275 |
| NCI-H209 | SCLC-A | 8075 |
| COR-L47 | SCLC-A | 10278 |
| NCI-H1436 | SCLC-A | 12795 |
| SW1271 | SCLC-Y | 25460 |
| NCI-H82 | SCLC-N | >30000 |
| NCI-H446 | SCLC-N | >30000 |
| NCI-H524 | SCLC-N | >30000 |
| NCI-H1836 | SCLC-A | >30000 |
| HCC-33 | SCLC-N | >30000 |

E

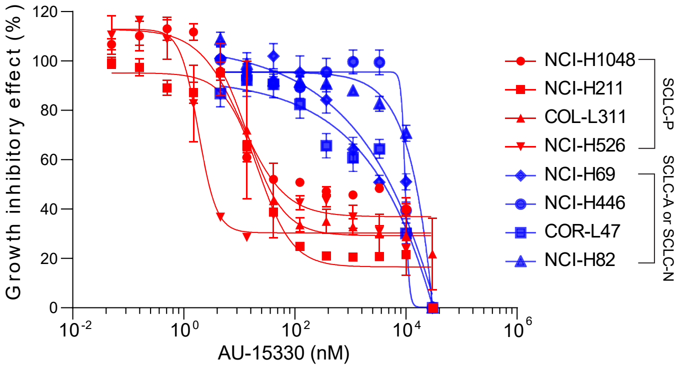

Figure S2

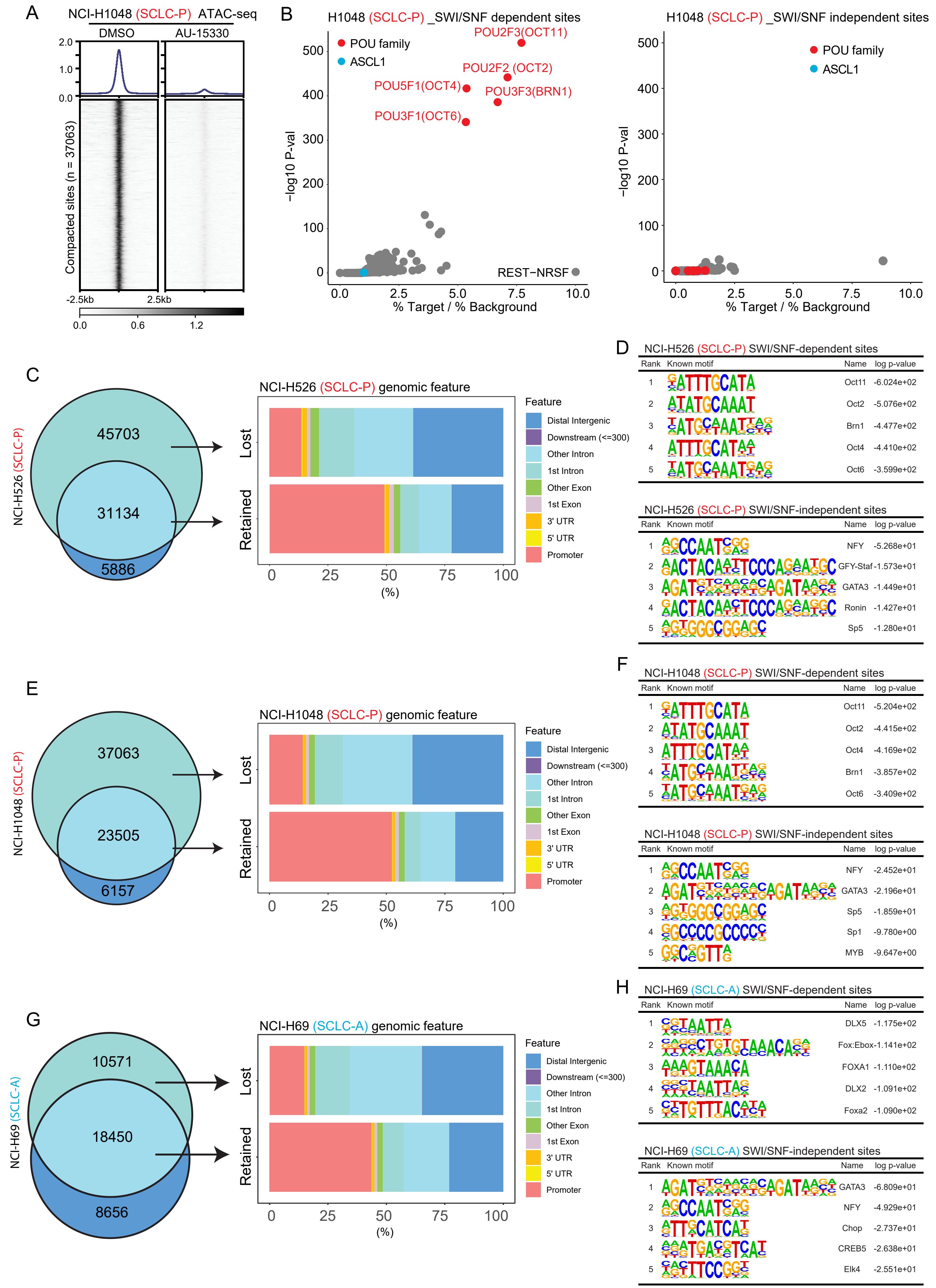
A
NCI-H1048 (SCLC-P) ATAC-seq
DMSO
AU-15330
2.0
1.0
0.0
Compacted sites (n = 37063)
-2.5kb
2.5kb
0.0
0.6
1.2
B
H1048 (SCLC-P) _SWI/SNF dependent sites
POU family
ASCL1
POU2F3(OCT11)
POU2F2 (OCT2)
POU5F1(OCT4)
POU3F3(BRN1)
POU3F1(OCT6)
REST−NRSF
−log10 P-val
% Target / % Background
H1048 (SCLC-P) _SWI/SNF independent sites
POU family
ASCL1
−log10 P-val
% Target / % Background
C
NCI-H526 (SCLC-P)
45703
31134
5886
NCI-H526 (SCLC-P) genomic feature
Lost
Retained
(%)
Feature
Distal Intergenic
Downstream (<=300)
Other Intron
1st Intron
Other Exon
1st Exon
3' UTR
5' UTR
Promoter
D
NCI-H526 (SCLC-P) SWI/SNF-dependent sites
Rank Known motif Name log p-value
1 Oct11 -6.024e+02
2 Oct2 -5.076e+02
3 Brn1 -4.477e+02
4 Oct4 -4.410e+02
5 Oct6 -3.599e+02
NCI-H526 (SCLC-P) SWI/SNF-independent sites
Rank Known motif Name log p-value
1 NFY -5.268e+01
2 GFY-Staf -1.573e+01
3 GATA3 -1.449e+01
4 Ronin -1.427e+01
5 Sp5 -1.280e+01
E
NCI-H1048 (SCLC-P)
37063
23505
6157
NCI-H1048 (SCLC-P) genomic feature
Lost
Retained
(%)
Feature
Distal Intergenic
Downstream (<=300)
Other Intron
1st Intron
Other Exon
1st Exon
3' UTR
5' UTR
Promoter
F
NCI-H1048 (SCLC-P) SWI/SNF-dependent sites
Rank Known motif Name log p-value
1 Oct11 -5.204e+02
2 Oct2 -4.415e+02
3 Oct4 -4.169e+02
4 Brn1 -3.857e+02
5 Oct6 -3.409e+02
NCI-H1048 (SCLC-P) SWI/SNF-independent sites
Rank Known motif Name log p-value
1 NFY -2.452e+01
2 GATA3 -2.196e+01
3 Sp5 -1.859e+01
4 Sp1 -9.780e+00
5 MYB -9.647e+00
G
NCI-H69 (SCLC-A)
10571
18450
8656
NCI-H69 (SCLC-A) genomic feature
Lost
Retained
(%)
Feature
Distal Intergenic
Downstream (<=300)
Other Intron
1st Intron
Other Exon
1st Exon
3' UTR
5' UTR
Promoter
H
NCI-H69 (SCLC-A) SWI/SNF-dependent sites
Rank Known motif Name log p-value
1 DLX5 -1.175e+02
2 Fox:Ebox -1.141e+02
3 FOXA1 -1.110e+02
4 DLX2 -1.091e+02
5 Foxa2 -1.090e+02
NCI-H69 (SCLC-A) SWI/SNF-independent sites
Rank Known motif Name log p-value
1 GATA3 -6.809e+01
2 NFY -4.929e+01
3 Chop -2.737e+01
4 CREB5 -2.638e+01
5 Elk4 -2.551e+01

### Figure S3

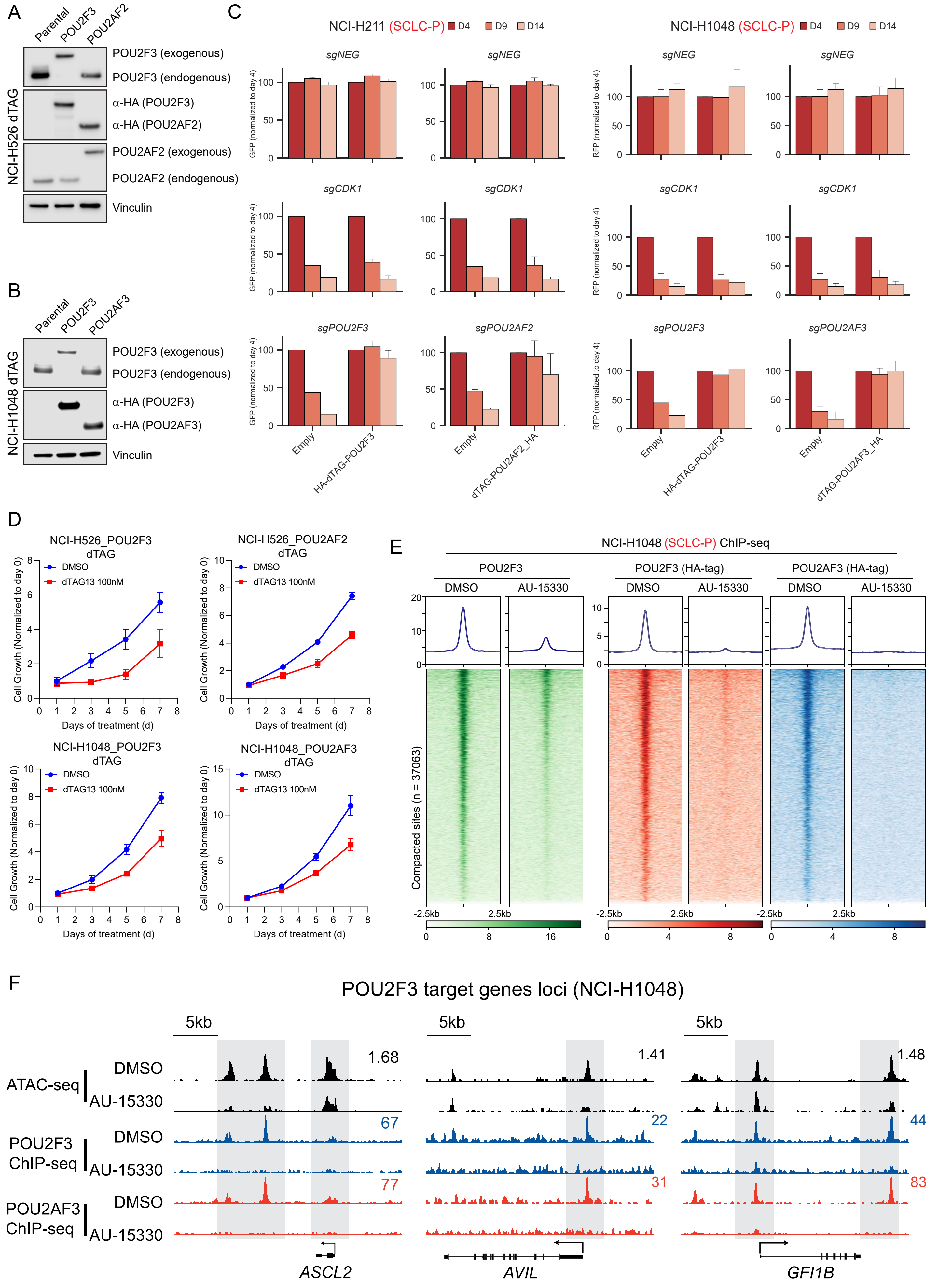

A
Parental
POU2F3
POU2AF2
NCI-H526 dTAG
POU2F3 (exogenous)
POU2F3 (endogenous)
α-HA (POU2F3)
α-HA (POU2AF2)
POU2AF2 (exogenous)
POU2AF2 (endogenous)
Vinculin
B
Parental
POU2F3
POU2AF3
NCI-H1048 dTAG
POU2F3 (exogenous)
POU2F3 (endogenous)
α-HA (POU2F3)
α-HA (POU2AF3)
Vinculin
C
NCI-H211 (SCLC-P) D4 D9 D14
NCI-H1048 (SCLC-P) D4 D9 D14
sgNEG
sgCDK1
sgPOU2F3
sgPOU2AF2
sgPOU2AF3
GFP (normalized to day 4)
RFP (normalized to day 4)
Empty
HA-dTAG-POU2F3
dTAG-POU2AF2_HA
dTAG-POU2AF3_HA
D
NCI-H526_POU2F3 dTAG
NCI-H526_POU2AF2 dTAG
NCI-H1048_POU2F3 dTAG
NCI-H1048_POU2AF3 dTAG
DMSO
dTAG13 100nM
Cell Growth (Normalized to day 0)
Days of treatment (d)
E
NCI-H1048 (SCLC-P) ChIP-seq
POU2F3
POU2F3 (HA-tag)
POU2AF3 (HA-tag)
DMSO
AU-15330
Compacted sites (n = 37063)
-2.5kb
2.5kb
F
POU2F3 target genes loci (NCI-H1048)
5kb
ATAC-seq
POU2F3 ChIP-seq
POU2AF3 ChIP-seq
DMSO
AU-15330
1.68
67
77
1.41
22
31
1.48
44
83
ASCL2
AVIL
GFI1B

### Figure S4

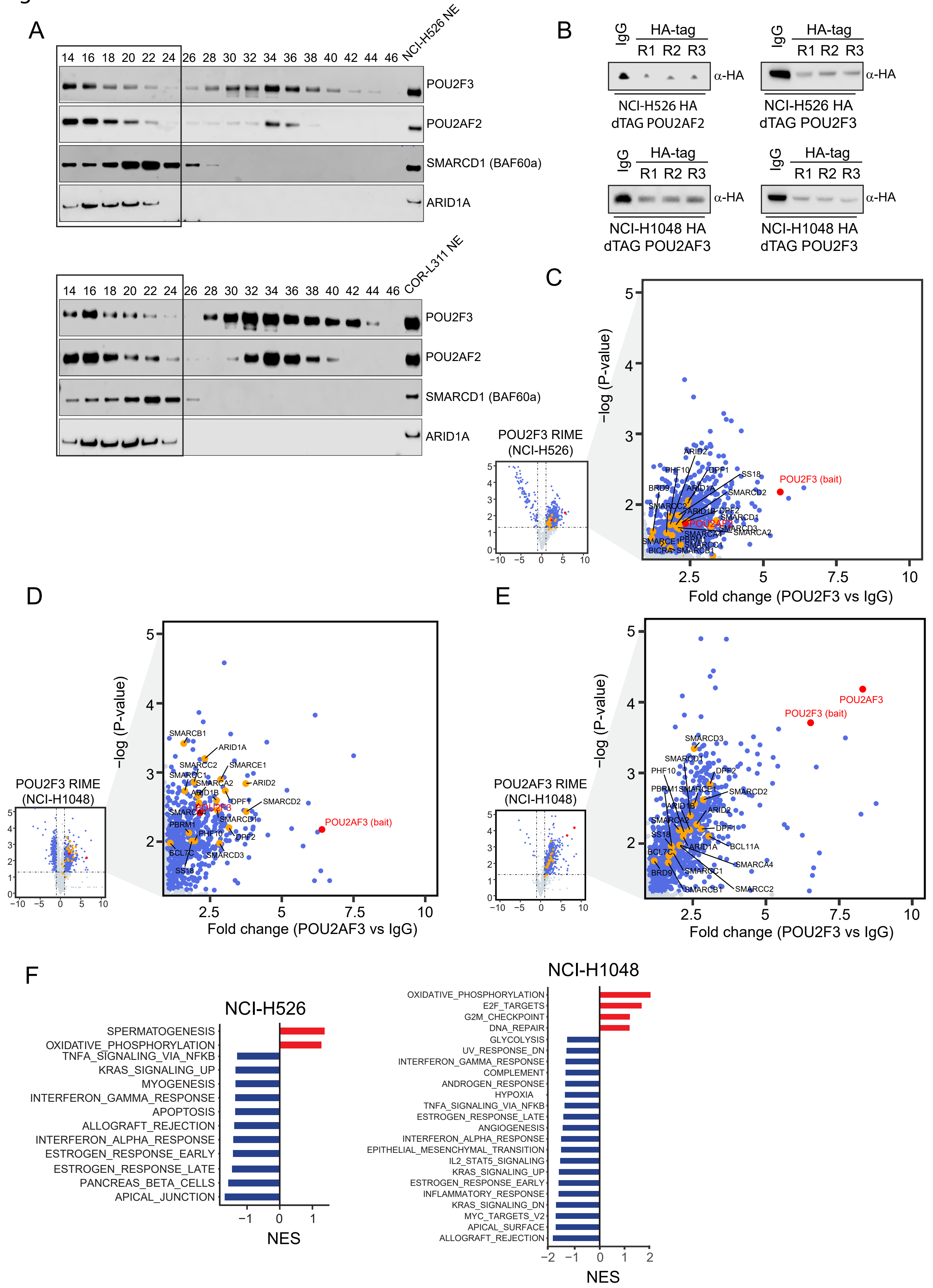

A
14 16 18 20 22 24 26 28 30 32 34 36 38 40 42 44 46 NCI-H526 NE
POU2F3
POU2AF2
SMARCD1 (BAF60a)
ARID1A
14 16 18 20 22 24 26 28 30 32 34 36 38 40 42 44 46 COR-L311 NE
POU2F3
POU2AF2
SMARCD1 (BAF60a)
ARID1A
B
IgG HA-tag R1 R2 R3 α-HA
NCI-H526 HA dTAG POU2AF2
NCI-H526 HA dTAG POU2F3
NCI-H1048 HA dTAG POU2AF3
NCI-H1048 HA dTAG POU2F3
C
POU2F3 RIME (NCI-H526)
POU2F3 (bait)
−log (P-value)
Fold change (POU2F3 vs IgG)
D
POU2F3 RIME (NCI-H1048)
POU2AF3 (bait)
−log (P-value)
Fold change (POU2AF3 vs IgG)
E
POU2AF3 RIME (NCI-H1048)
POU2AF3
POU2F3 (bait)
−log (P-value)
Fold change (POU2F3 vs IgG)
F
NCI-H526
SPERMATOGENESIS
OXIDATIVE_PHOSPHORYLATION
TNFA_SIGNALING_VIA_NFKB
KRAS_SIGNALING_UP
MYOGENESIS
INTERFERON_GAMMA_RESPONSE
APOPTOSIS
ALLOGRAFT_REJECTION
INTERFERON_ALPHA_RESPONSE
ESTROGEN_RESPONSE_EARLY
ESTROGEN_RESPONSE_LATE
PANCREAS_BETA_CELLS
APICAL_JUNCTION
NES
NCI-H1048
OXIDATIVE_PHOSPHORYLATION
E2F_TARGETS
G2M_CHECKPOINT
DNA_REPAIR
GLYCOLYSIS
UV_RESPONSE_DN
INTERFERON_GAMMA_RESPONSE
COMPLEMENT
ANDROGEN_RESPONSE
HYPOXIA
TNFA_SIGNALING_VIA_NFKB
ESTROGEN_RESPONSE_LATE
INTERFERON_ALPHA_RESPONSE
ANGIOGENESIS
EPITHELIAL_MESENCHYMAL_TRANSITION
IL2_STAT5_SIGNALING
KRAS_SIGNALING_UP
ESTROGEN_RESPONSE_EARLY
INFLAMMATORY_RESPONSE
KRAS_SIGNALING_DN
MYC_TARGETS_V2
APICAL_SURFACE
ALLOGRAFT_REJECTION
NES

Figure S5

A

| SCLC cell line | Subtype | AU-24118 (nM) |
|---|---|---|
| NCI-H526 | SCLC-P | 3.01 |
| NCI-H1048 | SCLC-P | 7.16 |
| NCI-H211 | SCLC-P | 5.48 |
| COR-L311 | SCLC-P | 20.59 |
| NCI-H1436 | SCLC-A | 2210 |
| COR-L47 | SCLC-A | 2442 |
| NCI-H1963 | SCLC-A | 2599 |
| NCI-H1836 | SCLC-A | 6588 |
| NCI-H209 | SCLC-A | 8060 |
| NCI-H446 | SCLC-N | 8605 |
| HCC-33 | SCLC-N | 8905 |
| NCI-H69 | SCLC-A | 10052 |
| NCI-H524 | SCLC-N | 12015 |
| NCI-H82 | SCLC-N | 13169 |
| SW1271 | SCLC-Y | >30000 |

B

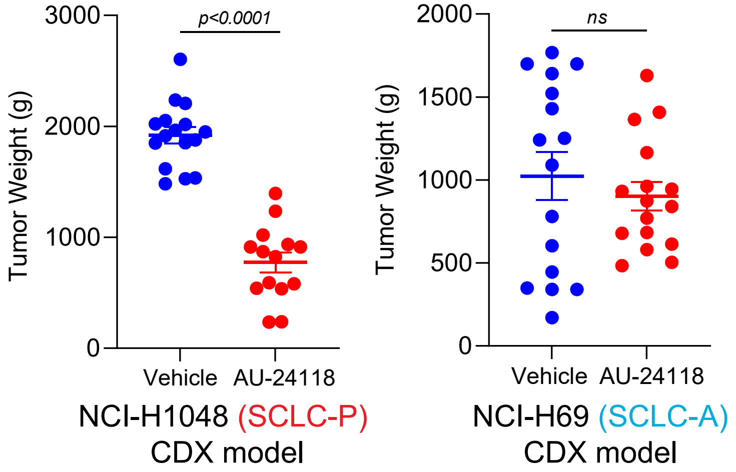


C

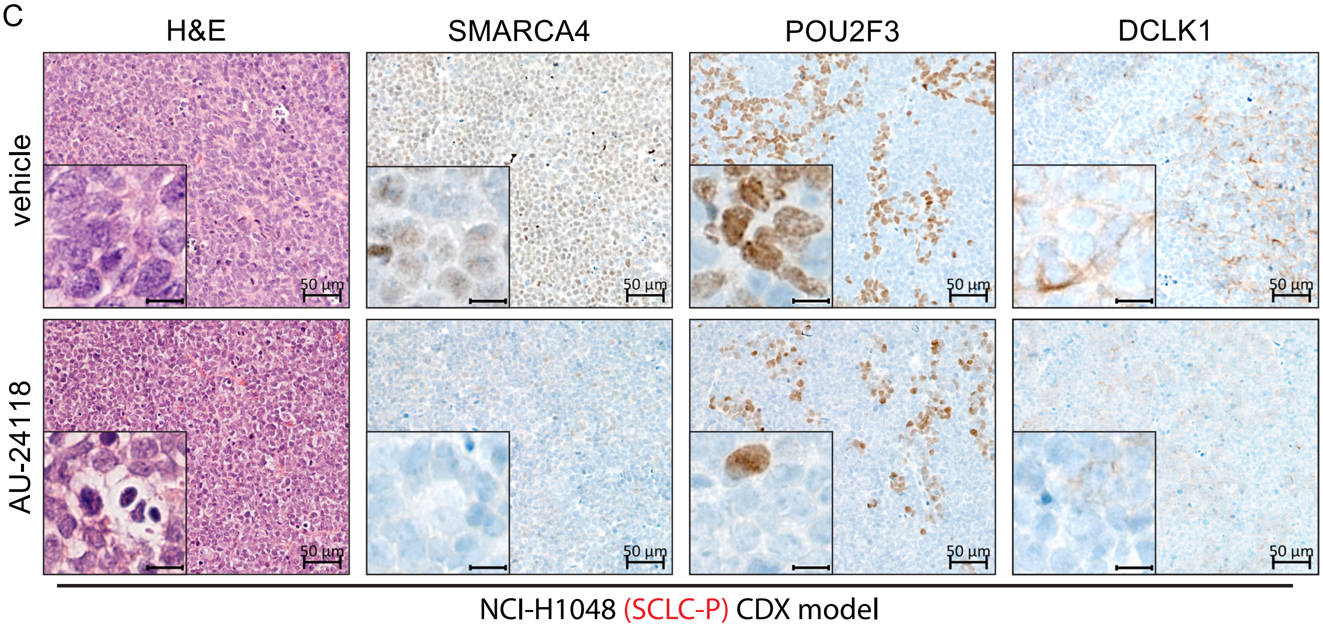


D

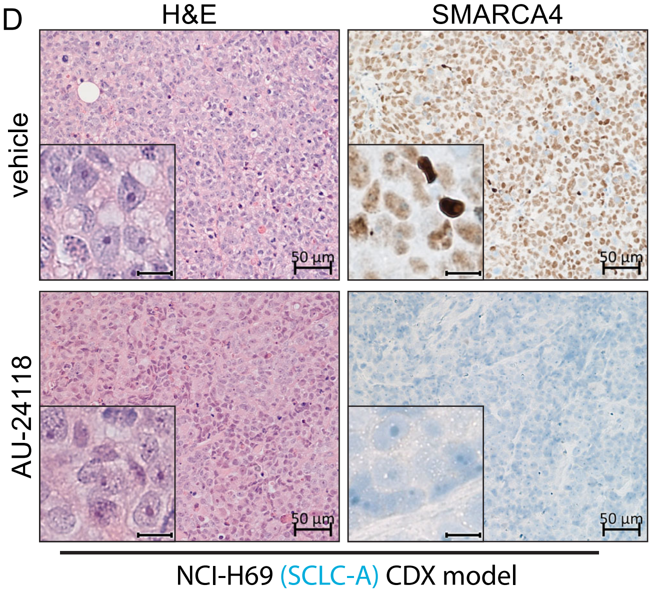

### Figure S6

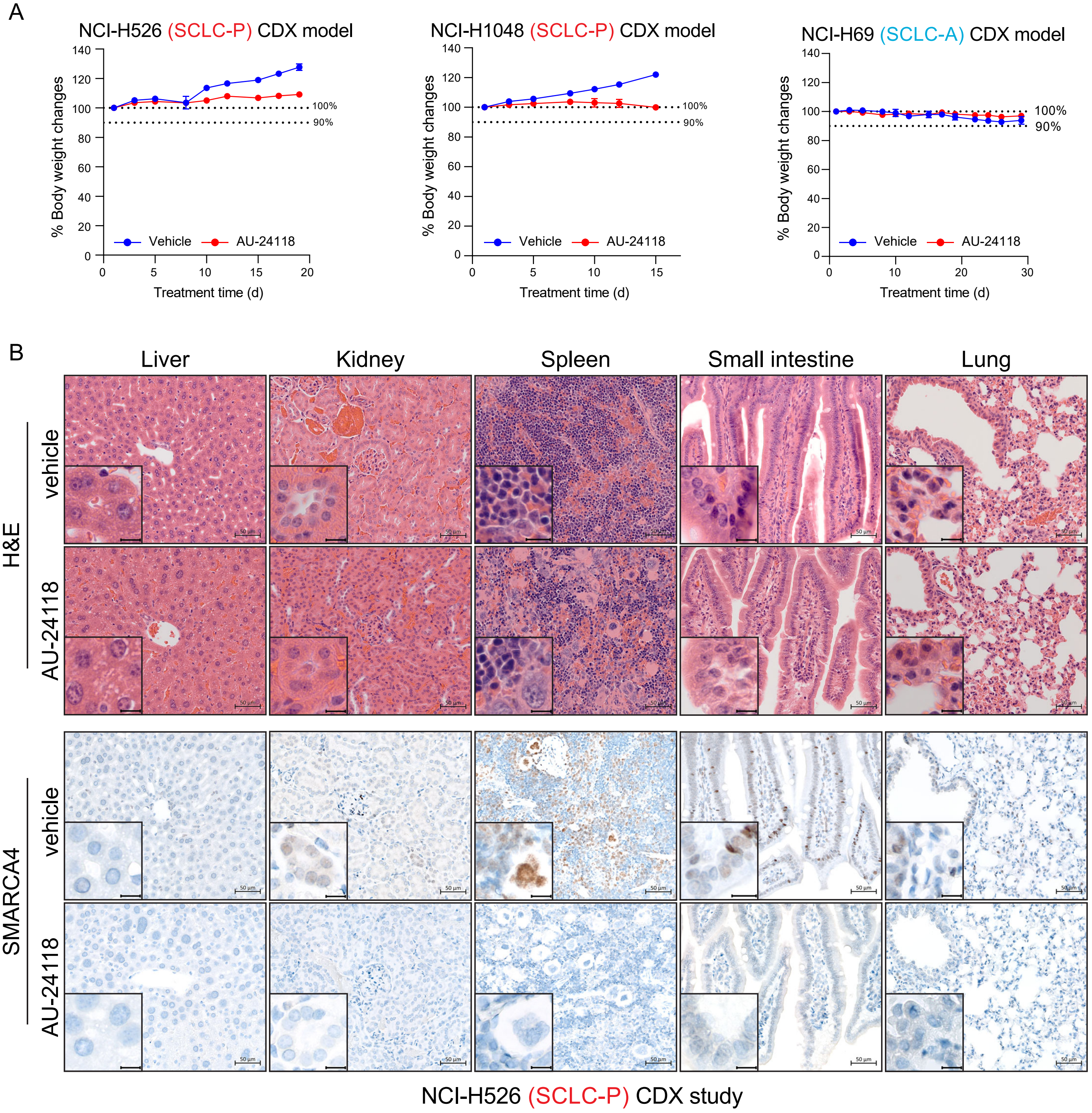
A
NCI-H526 (SCLC-P) CDX model
NCI-H1048 (SCLC-P) CDX model
NCI-H69 (SCLC-A) CDX model
% Body weight changes
Treatment time (d)
Vehicle
AU-24118
100%
90%
B
Liver
Kidney
Spleen
Small intestine
Lung
H&E
SMARCA4
vehicle
AU-24118
NCI-H526 (SCLC-P) CDX study

Figure S7

A

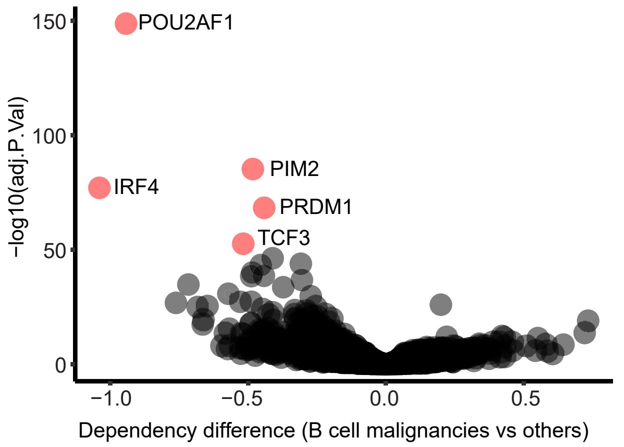


B

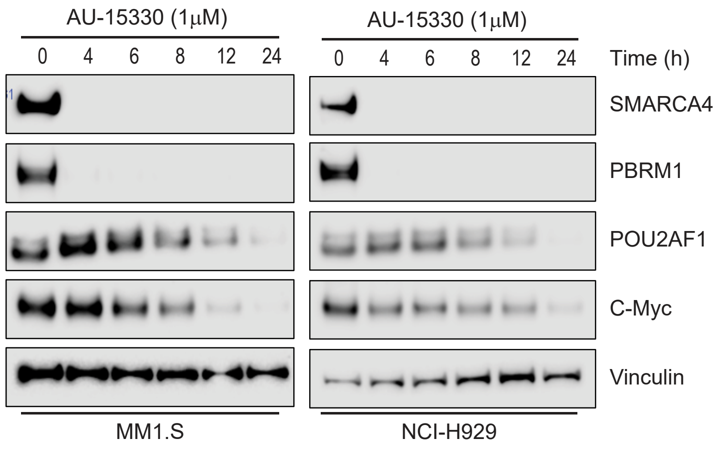


C

| MM cell line | AU-15330 (nM) |
|---|---|
| Karpas-25 | 2.5 |
| NCI-H929 | 5.02 |
| MM1.S | 9.74 |
| JIM1 | 20.76 |
| JIM3 | 32.16 |
| ANBL6 | 138.8 |
| Karpas-929 | 145.7 |
| Karpas-417 | 239.8 |
| RPMI-8226 | 470.6 |
| U266B1 | 1542 |

$IC_{50}$ summary of AU-15330 in MM cell lines

D

| DLBCL cell line | Subtype | AU-15330 (nM) |
|---|---|---|
| Karpas-422 | GCB | 22.83 |
| HT | GCB | 32.8 |
| Pfeiffer | GCB | 34.37 |
| SU-DHL-6 | GCB | 58.87 |
| DoHH2 | GCB | 73.45 |
| OCI-Ly1 | GCB | 164.4 |
| OCI-Ly7 | GCB | 172.5 |
| SU-DHL-5 | GCB | 306.5 |
| WSU-DLCL2 | GCB | 336.5 |
| DB | GCB | 363.6 |
| SU-DHL-4 | ABC | 370.1 |
| TMD8 | ABC | 370.9 |
| VAL | ABC | 439.1 |
| OCI-Ly3 | ABC | 496.9 |
| U2932 | ABC | 553.2 |
| RC-K8 | GCB or ABC | >30000 |

$IC_{50}$ summary of AU-15330 in DLBCL cell lines

### Figure S8

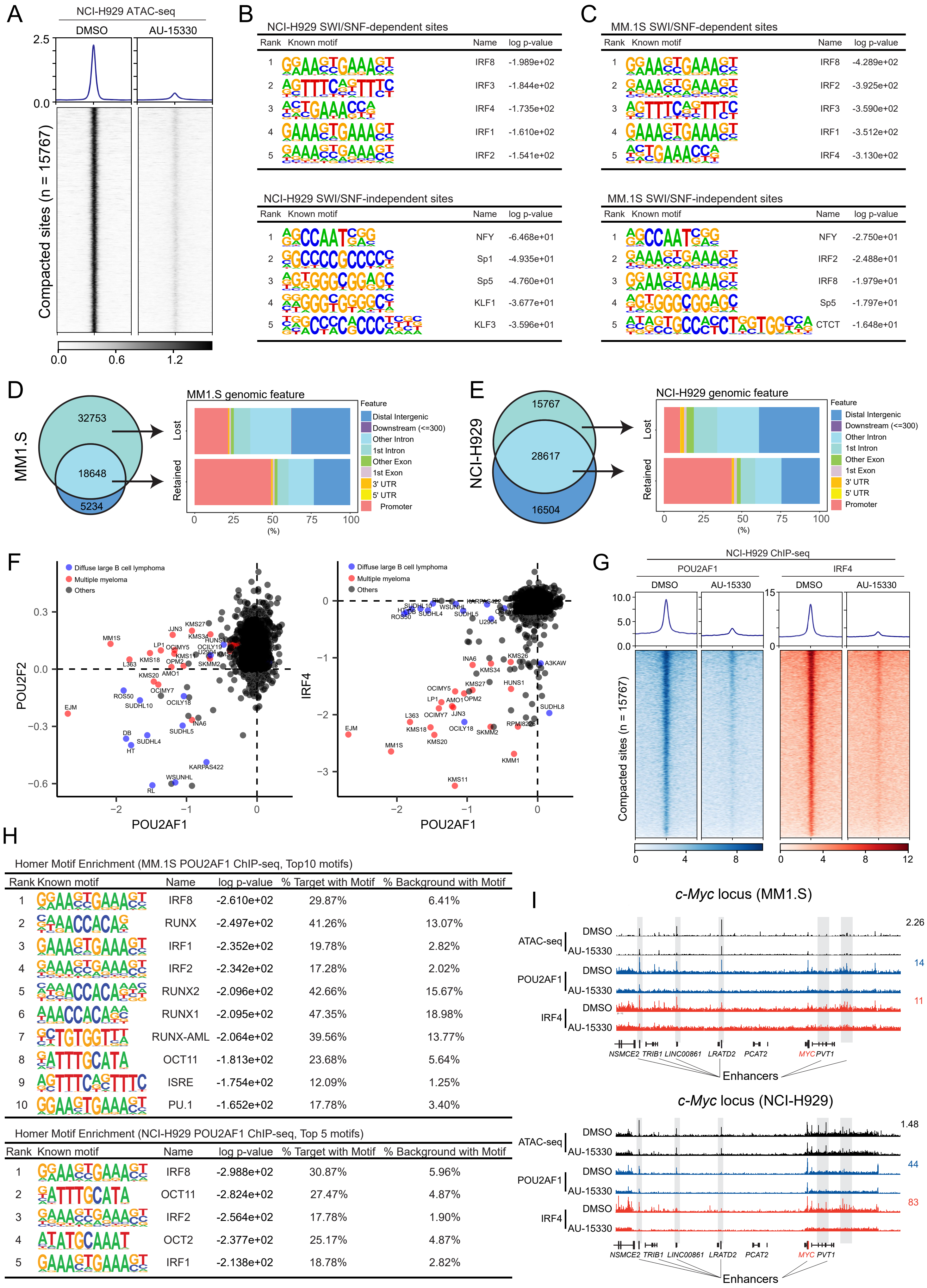
A
NCI-H929 ATAC-seq
DMSO
AU-15330
2.5
0.0
Compacted sites (n = 15767)
0.0
0.6
1.2
B
NCI-H929 SWI/SNF-dependent sites
Rank Known motif Name log p-value
1 IRF8 -1.989e+02
2 IRF3 -1.844e+02
3 IRF4 -1.735e+02
4 IRF1 -1.610e+02
5 IRF2 -1.541e+02
NCI-H929 SWI/SNF-independent sites
Rank Known motif Name log p-value
1 NFY -6.468e+01
2 Sp1 -4.935e+01
3 Sp5 -4.760e+01
4 KLF1 -3.677e+01
5 KLF3 -3.596e+01
C
MM.1S SWI/SNF-dependent sites
Rank Known motif Name log p-value
1 IRF8 -4.289e+02
2 IRF2 -3.925e+02
3 IRF3 -3.590e+02
4 IRF1 -3.512e+02
5 IRF4 -3.130e+02
MM.1S SWI/SNF-independent sites
Rank Known motif Name log p-value
1 NFY -2.750e+01
2 IRF2 -2.488e+01
3 IRF8 -1.979e+01
4 Sp5 -1.797e+01
5 CTCT -1.648e+01
D
MM1.S
32753
18648
5234
MM1.S genomic feature
Lost
Retained
Feature
Distal Intergenic
Downstream (<=300)
Other Intron
1st Intron
Other Exon
1st Exon
3' UTR
5' UTR
Promoter
0 25 50 75 100
(%)
E
NCI-H929
15767
28617
16504
NCI-H929 genomic feature
Lost
Retained
Feature
Distal Intergenic
Downstream (<=300)
Other Intron
1st Intron
Other Exon
1st Exon
3' UTR
5' UTR
Promoter
0 25 50 75 100
(%)
F
Diffuse large B cell lymphoma
Multiple myeloma
Others
POU2F2
POU2AF1
IRF4
POU2AF1
G
NCI-H929 ChIP-seq
POU2AF1
IRF4
DMSO
AU-15330
DMSO
AU-15330
Compacted sites (n = 15767)
0 4 8
0 4 8 12
H
Homer Motif Enrichment (MM.1S POU2AF1 ChIP-seq, Top10 motifs)
Rank Known motif Name log p-value % Target with Motif % Background with Motif
1 IRF8 -2.610e+02 29.87% 6.41%
2 RUNX -2.497e+02 41.26% 13.07%
3 IRF1 -2.352e+02 19.78% 2.82%
4 IRF2 -2.342e+02 17.28% 2.02%
5 RUNX2 -2.096e+02 42.66% 15.67%
6 RUNX1 -2.095e+02 47.35% 18.98%
7 RUNX-AML -2.064e+02 39.56% 13.77%
8 OCT11 -1.813e+02 23.68% 5.64%
9 ISRE -1.754e+02 12.09% 1.25%
10 PU.1 -1.652e+02 17.78% 3.40%
Homer Motif Enrichment (NCI-H929 POU2AF1 ChIP-seq, Top 5 motifs)
Rank Known motif Name log p-value % Target with Motif % Background with Motif
1 IRF8 -2.988e+02 30.87% 5.96%
2 OCT11 -2.824e+02 27.47% 4.87%
3 IRF2 -2.564e+02 17.78% 1.90%
4 OCT2 -2.377e+02 25.17% 4.87%
5 IRF1 -2.138e+02 18.78% 2.82%
I
c-Myc locus (MM1.S)
ATAC-seq
POU2AF1
IRF4
DMSO
AU-15330
2.26
14
11
NSMCE2 TRIB1 LINC00861 LRATD2 PCAT2 MYC PVT1
Enhancers
c-Myc locus (NCI-H929)
ATAC-seq
POU2AF1
IRF4
DMSO
AU-15330
1.48
44
83
NSMCE2 TRIB1 LINC00861 LRATD2 PCAT2 MYC PVT1
Enhancers

### Figure S9

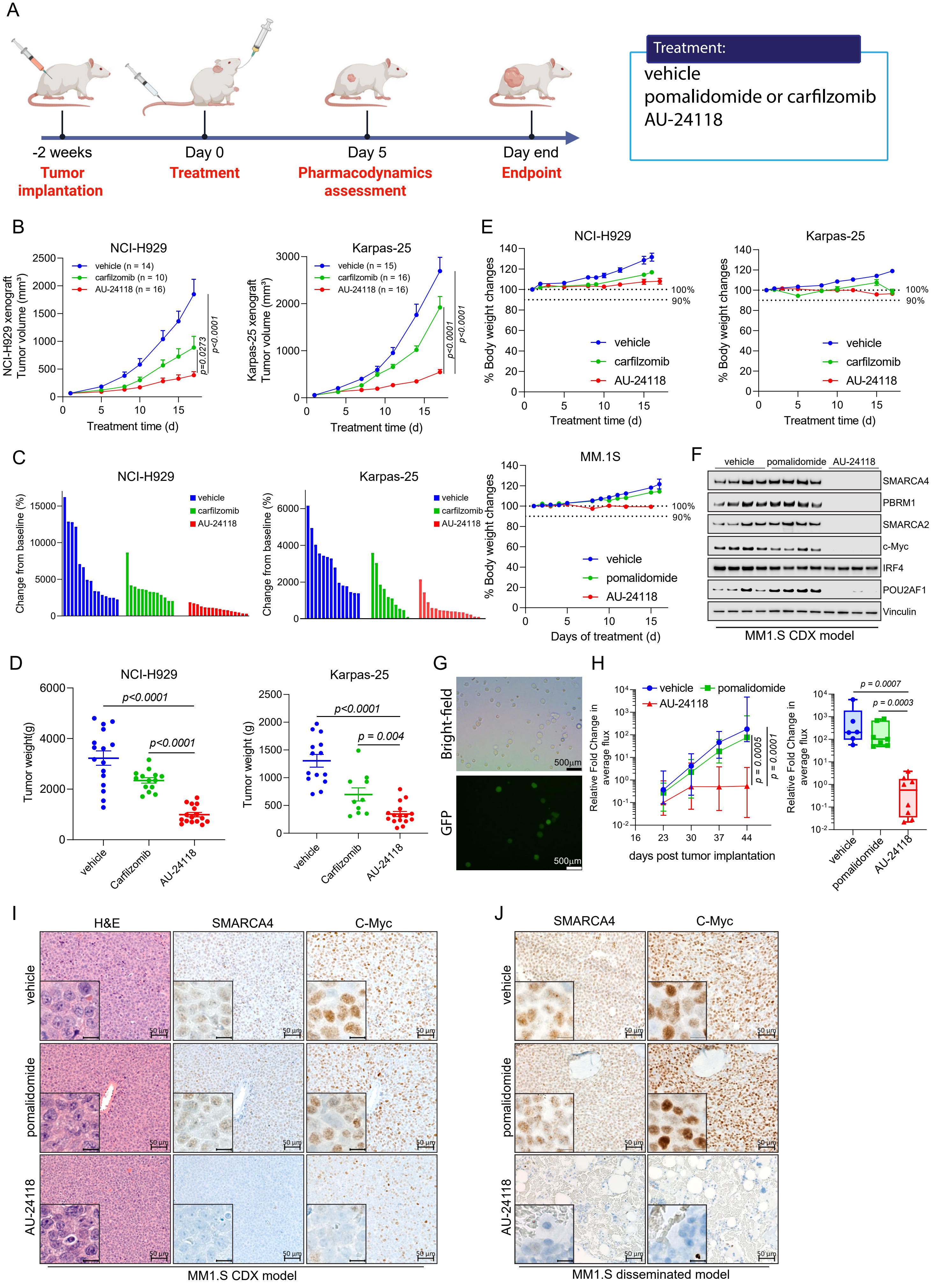

A
Treatment:
vehicle
pomalidomide or carfilzomib
AU-24118
-2 weeks
Tumor implantation
Day 0
Treatment
Day 5
Pharmacodynamics assessment
Day end
Endpoint
B
NCI-H929
vehicle (n = 14)
carfilzomib (n = 10)
AU-24118 (n = 16)
NCI-H929 xenograft Tumor volume (mm³)
Treatment time (d)
p=0.0273
p<0.0001
Karpas-25
vehicle (n = 15)
carfilzomib (n = 16)
AU-24118 (n = 16)
Karpas-25 xenograft Tumor volume (mm³)
p<0.0001
p<0.0001
C
NCI-H929
Karpas-25
vehicle
carfilzomib
AU-24118
Change from baseline (%)
D
NCI-H929
Karpas-25
Tumor weight(g)
Tumor weight (g)
p<0.0001
p = 0.004
vehicle
Carfilzomib
AU-24118
E
NCI-H929
Karpas-25
MM.1S
% Body weight changes
Treatment time (d)
Days of treatment (d)
100%
90%
vehicle
carfilzomib
pomalidomide
AU-24118
F
vehicle
pomalidomide
AU-24118
SMARCA4
PBRM1
SMARCA2
c-Myc
IRF4
POU2AF1
Vinculin
MM1.S CDX model
G
Bright-field
GFP
500μm
H
vehicle
pomalidomide
AU-24118
Relative Fold Change in average flux
days post tumor implantation
p = 0.0005
p = 0.0001
p = 0.0007
p = 0.0003
I
H&E
SMARCA4
C-Myc
vehicle
pomalidomide
AU-24118
50 μm
MM1.S CDX model
J
SMARCA4
C-Myc
MM1.S disseminated model

### Figure 1E

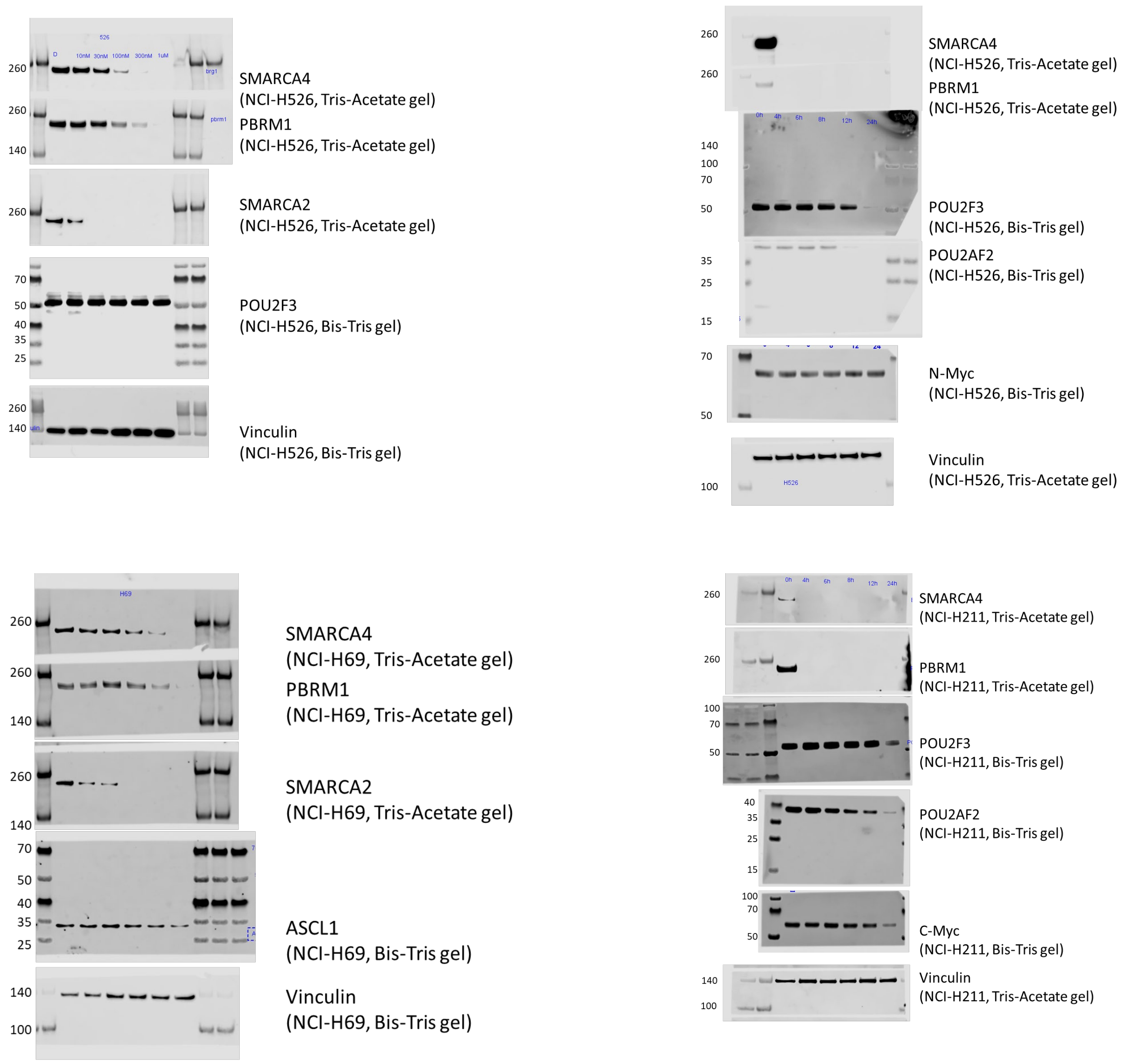

SMARCA4
(NCI-H526, Tris-Acetate gel)
PBRM1
(NCI-H526, Tris-Acetate gel)
SMARCA2
(NCI-H526, Tris-Acetate gel)
POU2F3
(NCI-H526, Bis-Tris gel)
Vinculin
(NCI-H526, Bis-Tris gel)
SMARCA4
(NCI-H526, Tris-Acetate gel)
PBRM1
(NCI-H526, Tris-Acetate gel)
POU2F3
(NCI-H526, Bis-Tris gel)
POU2AF2
(NCI-H526, Bis-Tris gel)
N-Myc
(NCI-H526, Bis-Tris gel)
Vinculin
(NCI-H526, Tris-Acetate gel)
SMARCA4
(NCI-H69, Tris-Acetate gel)
PBRM1
(NCI-H69, Tris-Acetate gel)
SMARCA2
(NCI-H69, Tris-Acetate gel)
ASCL1
(NCI-H69, Bis-Tris gel)
Vinculin
(NCI-H69, Bis-Tris gel)
SMARCA4
(NCI-H211, Tris-Acetate gel)
PBRM1
(NCI-H211, Tris-Acetate gel)
POU2F3
(NCI-H211, Bis-Tris gel)
POU2AF2
(NCI-H211, Bis-Tris gel)
C-Myc
(NCI-H211, Bis-Tris gel)
Vinculin
(NCI-H211, Tris-Acetate gel)

Fig S1C

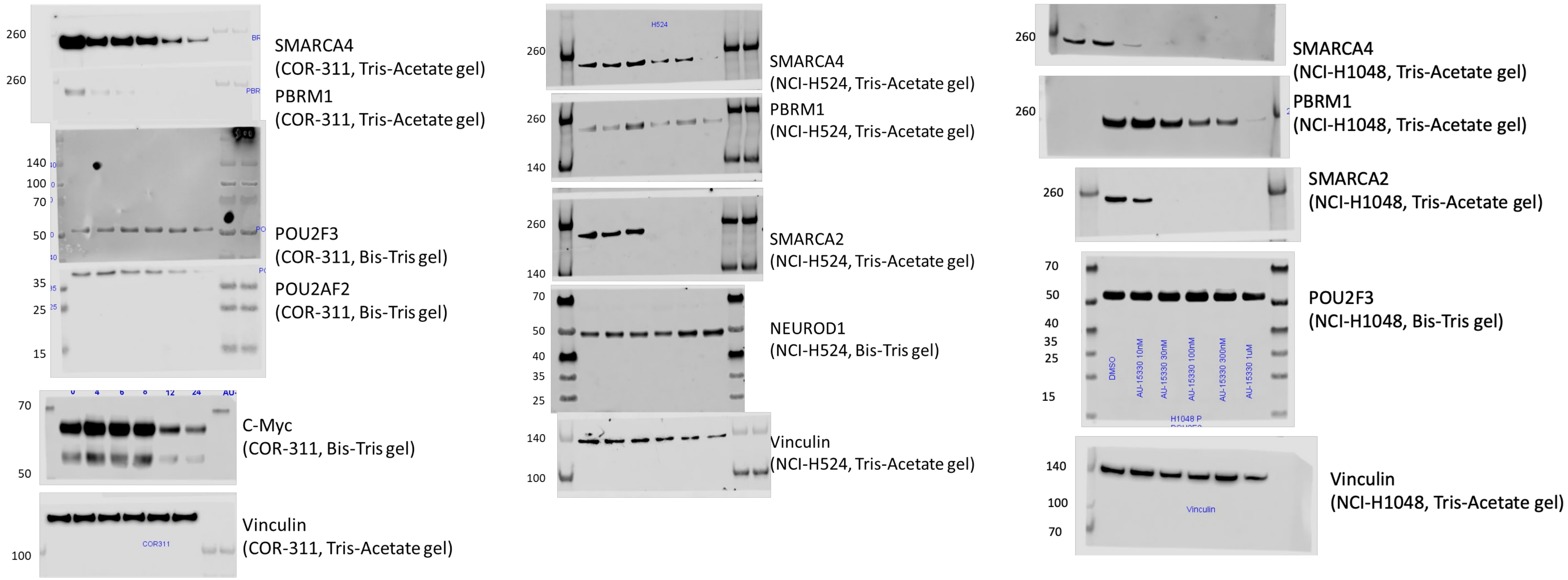

260
260
SMARCA4
(COR-311, Tris-Acetate gel)
PBRM1
(COR-311, Tris-Acetate gel)
140
100
70
50
POU2F3
(COR-311, Bis-Tris gel)
35
25
15
POU2AF2
(COR-311, Bis-Tris gel)
0 4 6 8 12 24
70
50
C-Myc
(COR-311, Bis-Tris gel)
100
COR311
Vinculin
(COR-311, Tris-Acetate gel)
H524
260
SMARCA4
(NCI-H524, Tris-Acetate gel)
260
140
PBRM1
(NCI-H524, Tris-Acetate gel)
260
140
SMARCA2
(NCI-H524, Tris-Acetate gel)
70
50
40
35
25
NEUROD1
(NCI-H524, Bis-Tris gel)
140
100
Vinculin
(NCI-H524, Tris-Acetate gel)
260
SMARCA4
(NCI-H1048, Tris-Acetate gel)
260
PBRM1
(NCI-H1048, Tris-Acetate gel)
260
SMARCA2
(NCI-H1048, Tris-Acetate gel)
70
50
40
35
25
15
DMSO
AU-15330 10nM
AU-15330 30nM
AU-15330 100nM
AU-15330 300nM
AU-15330 1uM
H1048 P
POU2F3
(NCI-H1048, Bis-Tris gel)
140
100
70
Vinculin
Vinculin
(NCI-H1048, Tris-Acetate gel)

### Fig 3C

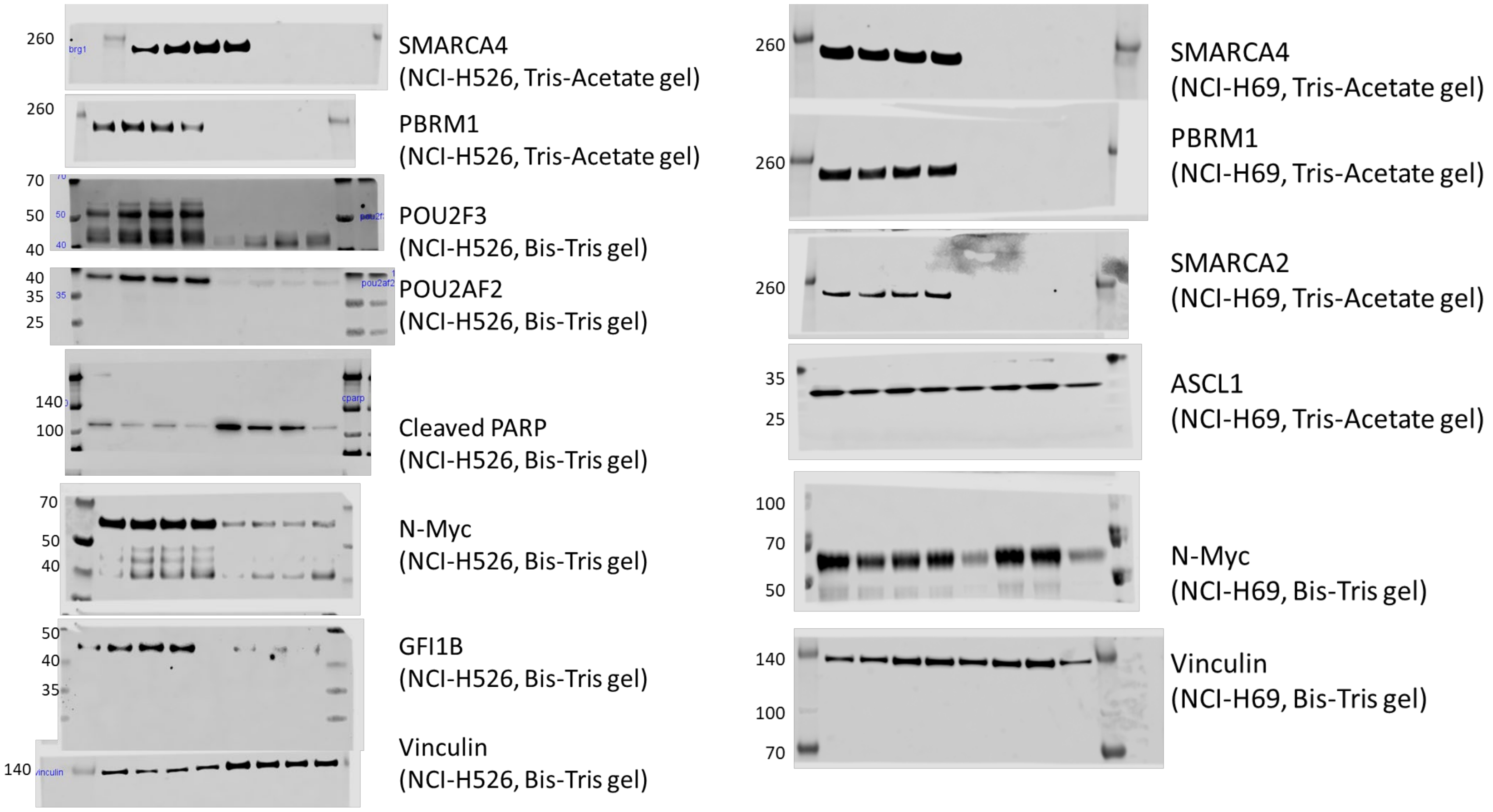

260
SMARCA4
(NCI-H526, Tris-Acetate gel)
260
PBRM1
(NCI-H526, Tris-Acetate gel)
70
50
40
POU2F3
(NCI-H526, Bis-Tris gel)
40
35
25
POU2AF2
(NCI-H526, Bis-Tris gel)
140
100
Cleaved PARP
(NCI-H526, Bis-Tris gel)
70
50
40
N-Myc
(NCI-H526, Bis-Tris gel)
50
40
35
GFI1B
(NCI-H526, Bis-Tris gel)
140
Vinculin
(NCI-H526, Bis-Tris gel)
260
SMARCA4
(NCI-H69, Tris-Acetate gel)
260
PBRM1
(NCI-H69, Tris-Acetate gel)
260
SMARCA2
(NCI-H69, Tris-Acetate gel)
35
25
ASCL1
(NCI-H69, Tris-Acetate gel)
100
70
50
N-Myc
(NCI-H69, Bis-Tris gel)
140
100
70
Vinculin
(NCI-H69, Bis-Tris gel)

#### Fig S3A

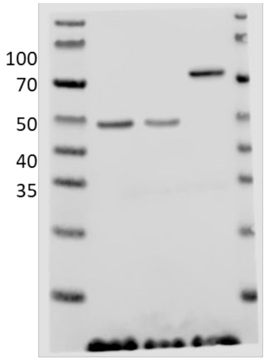


POU2AF2
(NCI-H526, dTAG-HA Bis-Tris gel)

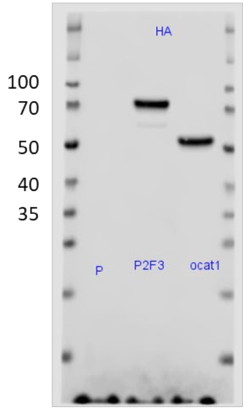


HA-tag
(NCI-H526, dTAG-HA Bis-Tris gel)

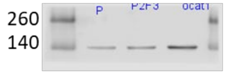


Vinculin
(NCI-H526, dTAG-HA Bis-Tris gel)

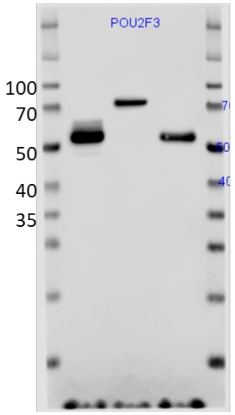


POU2F3
(NCI-H526, dTAG-HA Bis-Tris gel)

#### Fig S3B

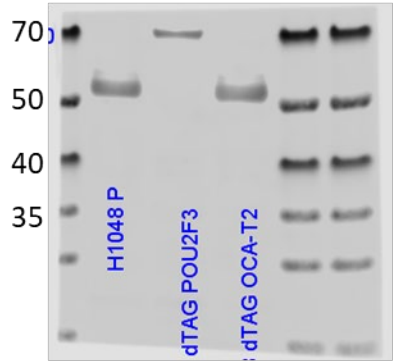


POU2F3
(NCI-H1048, dTAG-HA Bis-Tris gel)

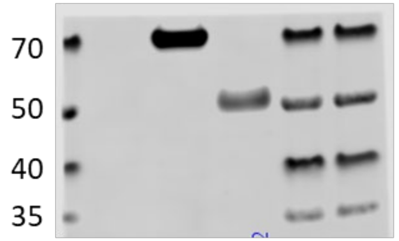


HA-tag
(NCI-H1048, dTAG-HA Bis-Tris gel)

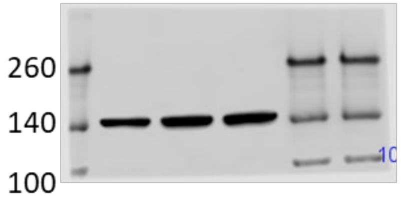


Vinculin
(NCI-H1048, Bis-Tris gel)

#### Fig S4A (Upper)

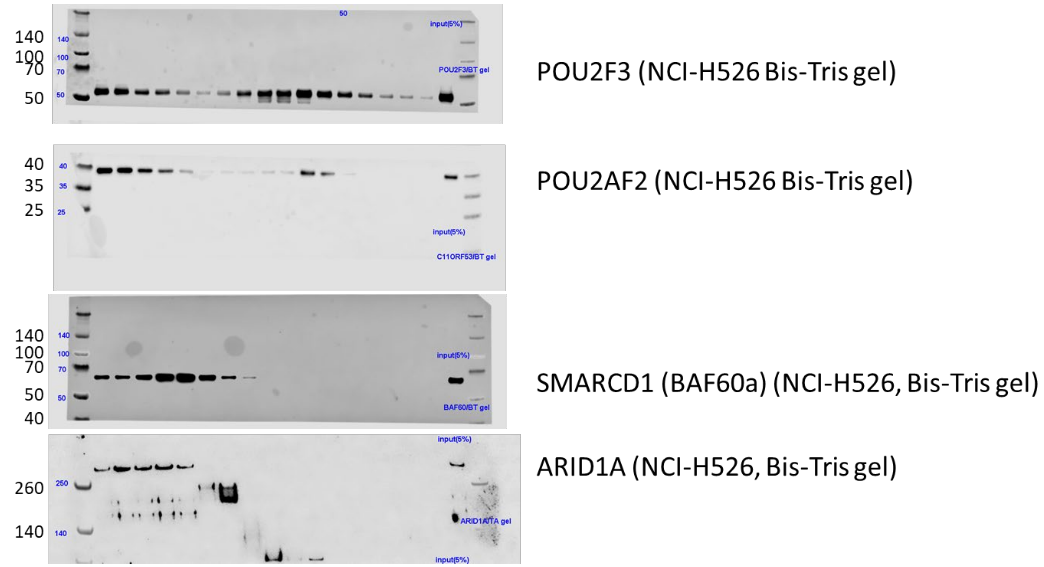


#### Fig S4A (Bottom)

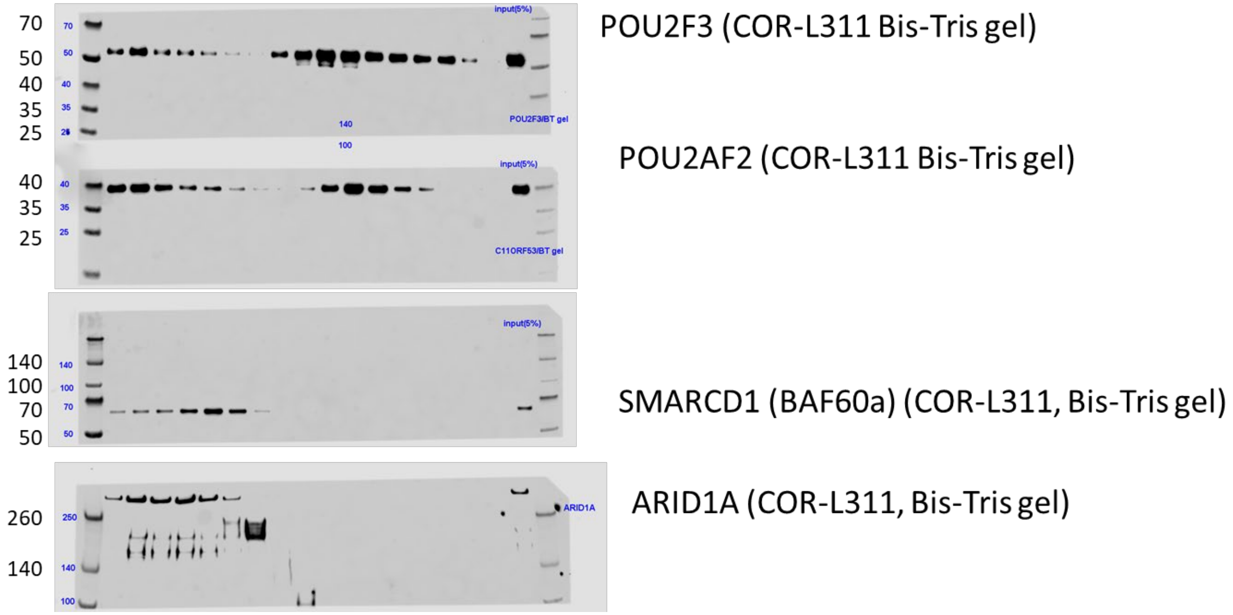


#### Fig S4B

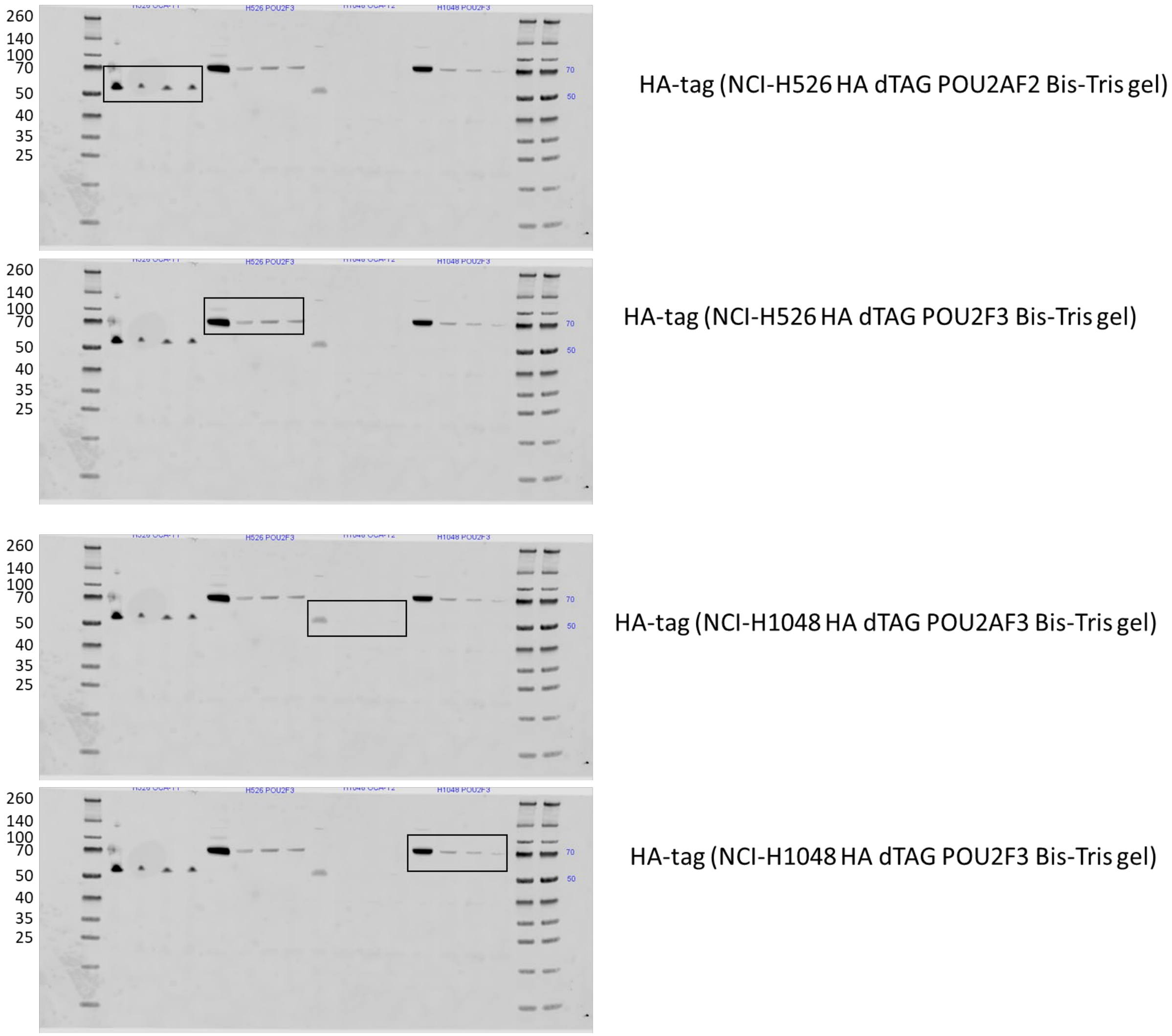

#### Fig S7B left

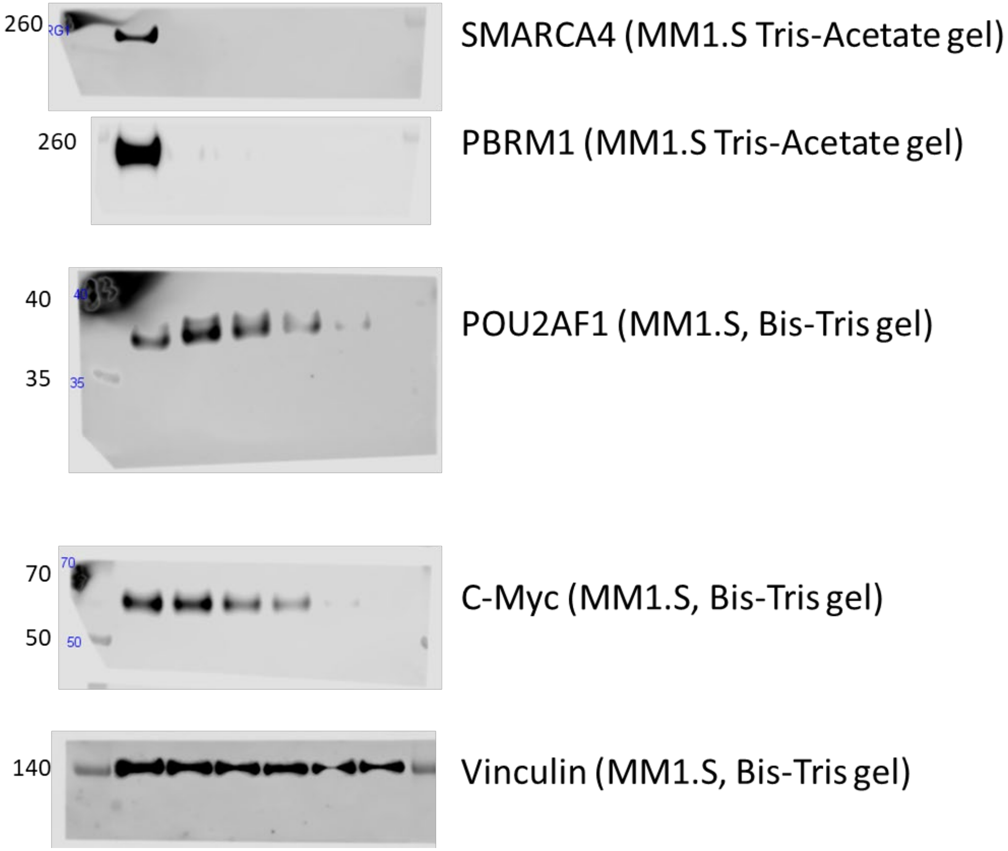


#### Fig S7B right

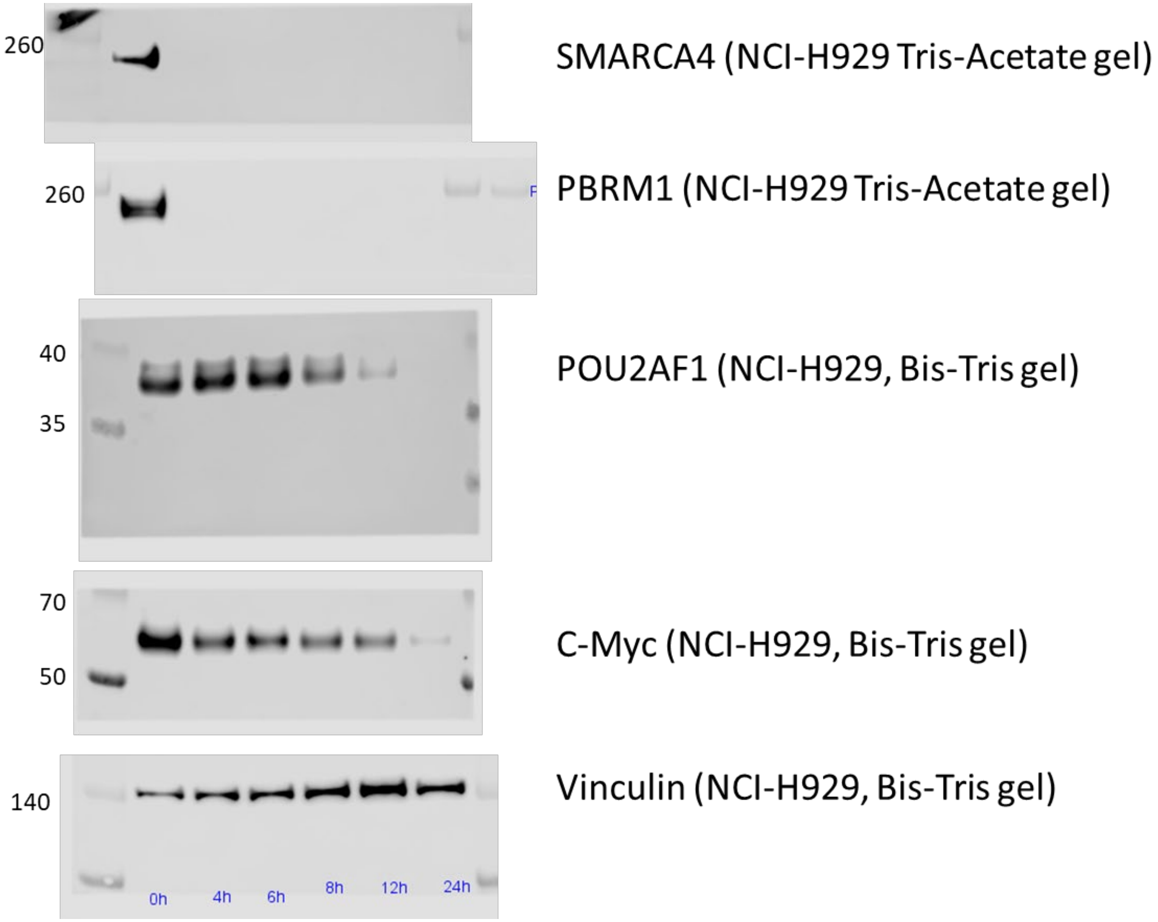

### Fig S9F

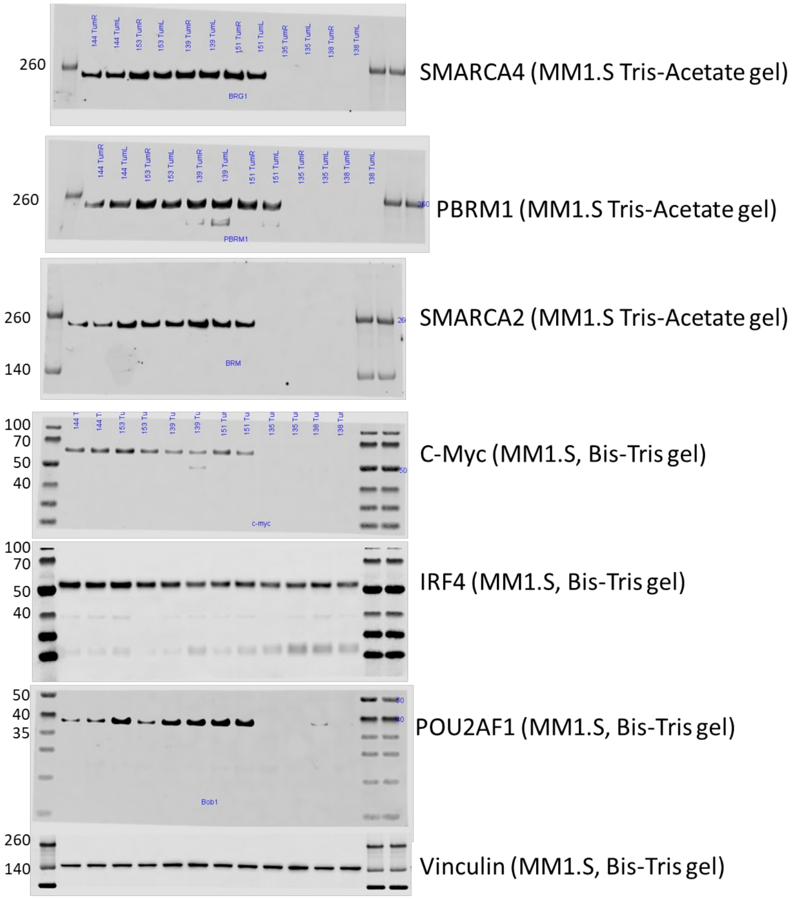
260
SMARCA4 (MM1.S Tris-Acetate gel)
260
PBRM1 (MM1.S Tris-Acetate gel)
260
140
SMARCA2 (MM1.S Tris-Acetate gel)
100
70
50
40
C-Myc (MM1.S, Bis-Tris gel)
100
70
50
40
IRF4 (MM1.S, Bis-Tris gel)
50
40
35
POU2AF1 (MM1.S, Bis-Tris gel)
260
140
Vinculin (MM1.S, Bis-Tris gel)
